# Dynamin forms liquid-like condensates at synapses to support ultrafast endocytosis

**DOI:** 10.1101/2022.06.01.494432

**Authors:** Yuuta Imoto, Ye Ma, Kie Itoh, Eva-Maria Blumrich, Hideaki T. Matsubayashi, Jian Liu, Bin Wu, Michael A. Cousin, Taekjip Ha, Takanari Inoue, Shigeki Watanabe

**Author notes:** Correspondence to: Shigeki Watanabe, and Yuuta Imoto.

## Abstract

Endocytosis at synapses is accelerated by the pre-accumulation of Dynamin 1xA at the endocytic zone by Syndapin 1. However, it is unclear how these proteins support the ultrafast kinetics of endocytosis. Here we report that these proteins phase separate at the presynaptic endocytic zone where ultrafast endocytosis takes place. Specifically, the proline-rich motif of Dynamin 1xA interacts with the Src-Homology 3 domain of Syndapin 1 and forms liquid-like condensates. Single-particle tracking of Dynamin 1xA molecules at synapses shows that their diffusion slows down substantially when they are in the condensates, indicating the presence of molecular crowding and intermolecular interaction. When Dynamin 1xA is mutated to disrupt its interaction with Syndapin 1 the condensates do not form. Thus, the liquid-like assembly of these endocytic proteins provides a catalytic platform for ultrafast endocytosis.

## Introduction

Phase separation can accumulate proteins to certain areas of a cell and compartmentalize cellular functions (Hyman et al., 2014; Li et al., 2012; Shin and Brangwynne, 2017). Proteins that partition into the same phase are typically multivalent and have a weak hydrophobic interaction with each other (Hyman et al., 2014). When the critical concentrations are reached, these proteins partition into high- and low-concentration phases to minimize thermodynamic energy (Hyman et al., 2014). Recent studies suggest that this principle governs presynaptic organization (Hayashi et al., 2021; Milovanovic and De Camilli, 2017). Synapsin 1 forms condensates through its intrinsically disordered region to form the synaptic vesicle cluster (Milovanovic et al., 2018; Pechstein et al., 2020), potentially via its interaction with synaptophysin (Park et al., 2021). Likewise, active zone proteins also form distinct molecular condensates (Wu et al., 2019, 2021) and recruit synaptic vesicles near calcium channels (Wu et al., 2021). Condensate formation at the active zone is regulated developmentally (McDonald et al., 2020) and by activity (Emperador-Melero et al., 2021), thereby allowing dynamic exchange of materials including synaptic vesicles between phases throughout the lifetime of synapses.

Like the exocytic machinery, endocytic proteins that control the recycling of synaptic vesicles also possess the molecular characteristics necessary for phase separation and colocalize at synapses. A splice variant of Dynamin 1, Dyn1xA, is localized at the presynaptic endocytic zone by its interaction with the Fer-CIP4 homology-BAR (F-BAR) containing protein Syndapin 1 (Imoto et al., 2021). The C-terminus of Dyn1xA contains a 20 amino acid extension with extra binding motifs for SH3-containing proteins (Huang et al., 2004), providing multivalency. This region is essential for ultrafast endocytosis – another splice variant, Dyn1xB, which lacks this extension, cannot support ultrafast endocytosis (Imoto et al., 2021). Like other proteins that phase separate (Banani et al., 2017; Li et al., 2012; Milovanovic et al., 2018; Pechstein et al., 2020), this interaction is mediated by low-complexity motifs: the proline-rich motif (PRM) of Dyn1xA and the Src-Homology 3 (SH3) domain of Syndapin 1. This interaction is tightly controlled by the phosphorylation status of Dyn1xA PRM (Anggono et al., 2006), another trend among low-complexity motifs that form condensates. All these factors point to the possibility that Dyn1xA and Syndapin 1 may phase separate at synapses.

However, both dynamin and syndapin form rigid scaffolding structures on the membrane during endocytosis (Daumke et al., 2014; Haucke et al., 2011). These structures are typically formed on the highly curved membrane around the neck of endocytic pits (Daumke et al., 2014; Haucke et al., 2011; Rao et al., 2010; Taylor et al., 2012). However, cryo-electron microscopy studies of other F-BAR proteins (Frost et al., 2008) and molecular dynamics simulations (Mahmood et al., 2019) suggest that Syndapin 1 can also bind to flat membranes. Likewise, Dyn1 can localize to the scaffold-like clathrin lattice on a flat plasma membrane (Damke et al., 1994) and thereby accelerate clathrin-mediated endocytosis (Srinivasan et al., 2018). Furthermore, Dyn1 can form linear oligomers (Faelber et al., 2011) and accumulate on flat membranes by itself (Taylor et al., 2012). Thus, Dyn1xA and Syndapin 1 could pre-accumulate at the endocytic zone either via liquid-like condensates or scaffolding complexes.

In this study, we performed phase separation assays with purified recombinant proteins as well as exogenously expressed proteins in COS-7 cells and mouse hippocampal neurons. *In vitro* and in COS-7 cells, Dyn1xA and Syndapin 1 formed liquid-like droplets. When the phosphomimetic form of Dyn1xA (S774/778D) (Anggono et al., 2006) was used in these assays, Dyn1xA did not form liquid-like condensates. Likewise, in neurons, wild-type Dyn1xA also exhibited liquid-like behaviors: fluorescence recovered after photo-bleaching and the droplets dispersed when aliphatic alcohol was applied. To further test whether Dyn1xA forms liquid-like condensates in neurons, we performed a single-particle tracking assay at synapses and demonstrated the presence of a phase boundary on each punctum in intact synapses – the movement of molecules slows down once they enter the condensates. The phosphomimetic form of Dyn1xA cannot form liquid-like droplets and is diffuse throughout the cytoplasm similar to Syndapin 1 knock down phenotype which slows down ultrafast endocytosis in previous study (Imoto et al., 2021). Together, these results suggest that the liquid-like properties of Dyn1xA and Syndapin 1 are important for ultrafast endocytosis.

## Results

### Dyn1xA and Syndapin 1 form liquid-like condensates *in vitro*

Molecular condensation requires intrinsically-disordered regions as well as multivalent interactions with other proteins, and is often controlled by posttranslational modifications (Hyman et al., 2014; Milovanovic et al., 2018). The proline-rich motif of Dyn1xA and the SH3 domain of Syndapin 1 satisfy all these requirements: the Dyn1 PRM is highly disordered (Figure S1A) and binds Syndapin 1, a brain-specific isoform of syndapin (Anggono et al., 2006) in a manner dependent on dephosphorylation of the PRM. To test whether Dyn1xA molecules form condensates with Syndapin 1, we performed *in vitro* assays using purified recombinant full-length Dyn1xA and Syndapin 1 (Figure S1B), labelled with Alexa 488 and 549, respectively (see STAR methods). Interestingly, Dyn1xA alone seemed to form small aggregate-like structures (Figure S1C), while Syndapin 1 formed liquid droplet-like structures in the presence of polyethylene glycol (PEG) at physiological protein concentration and ionic strength (Figure S1C-H). When Dyn1xA and Syndapin 1 were mixed in the presence of polyethylene glycol (PEG) (∼2 µM and ∼20 µM, respectively), they coalesced into droplets (Figure 1A, Figure S1C). These droplets exhibit liquid-like properties such as spherical shapes (Figure 1A), fusion upon contact (Figure 1B), rapid re-rounding after fusion (Figure 1B), and rearrangement of molecules within puncta (Figure 1C, D). Of note, the kinetics of fluorescence recovery after photobleaching (FRAP) were dependent on the radius of photobleaching (Figure 1C-E & S1I; 900 nm; 30.0 s, 300 nm; 6.27 s), suggesting that the molecular rearrangement is dominated by protein diffusion rather than binding and unbinding (McSwiggen et al., 2019a).

**Figure 1.**
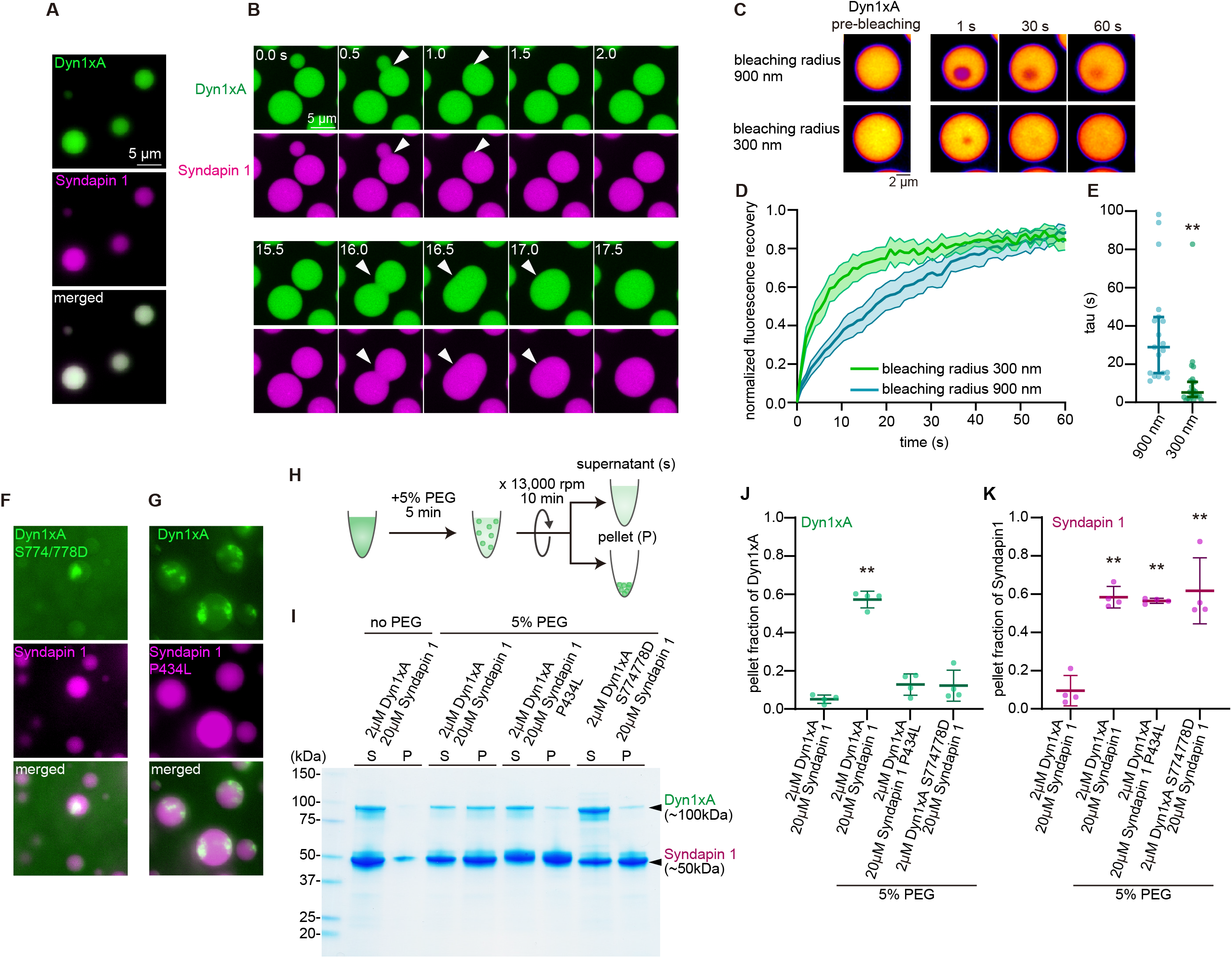
Dyn1xA exhibits liquid-like behaviors in *in vitro*. (A) Example images of purified Dyn1xA (labeled with Alexa488) and Syndapin1 (labeled with Alexa 549) under physiological salt concentration and with 10 % PEG. (B) Example time-lapse images of Dyn1xA and Syndapin1 droplets undergoing fusion. (C) Example time-lapse images of FRAP experiments in *in vitro*. Time indicates after the photobleaching. Dyn1xA and Syndapin1 droplets were photobleached at 480 nm using the region-of-interest (ROI) radius of 900 nm or 300 nm. (D) Normalized fluorescence recovery of Dyn1xA signals in different ROI sizes. Fluorescence signals were normalized between immediately (0 s) and 60 s after the photobleaching. Times indicate after the photobleaching. The median and 95% confidential interval are shown. (E) The recovery time constant of Dyn1xA signals following the photobleaching using different ROI sizes. The median and 95% confidence interval are shown. Each dot represents a Dyn1xA-Syndapin 1 droplet. p < 0.0001. Mann-Whitney test. (F) Example images of purified Dyn1xA S774/778D (labeled with Alexa488) and Syndapin1 (labeled with Alexa 549) under physiological salt concentration and with 10 % PEG. (G) Example images of purified Dyn1xA (labeled with Alexa488) and Syndapin1 P434L (labeled with Alexa 549) under physiological salt concentration and with 10 % PEG. All data are examined by n > 15 droplets from two independent protein purifications. (H) Schematics represents low-speed sedimentation assay. (I) Dyn1xA (2 μM) and Syndapin 1 (10 μM) were used with and without 10 % PEG. The supernatant (S) and pellet (P) were collected by centrifugation at ×13,000 rpm for 10 min and then proteins were subjected to SDS–PAGE and Gel code staining (I). (J, K) The pellet fraction of Dyn1xA (I) and Syndapin1 (J) were calculated from protein bands in supernatants and pellet fraction in (H). *, P < 0.05. **, P < 0.001. Mean ± SEM are shown. n = 4 from two independent protein purification.

To test if such coalescence is regulated by the weak hydrophobic PRM-SH3 interaction, we disrupted PRM-SH3 binding by introducing phosphomimetic mutations at S774 and S778 within the PRM of Dyn1xA (Anggono et al., 2006) (Xue et al., 2011) (Figure S1J, K). When recombinant Dyn1xA S774/778D and Syndapin 1 were mixed, Dyn1xA-S774/778D aggregated around the periphery of Syndapin 1 condensates (Figure 1F, G). Furthermore, sedimentation assays (Figure 1H) showed that only a small fraction of Dyn1xA S774/778D stays in the pellet (Figure 1I-K; 12.7 ± 5.6 %), with the rest in the soluble fraction (Figure 1I-K). The reciprocal experiment, mutating the SH3 domain of Syndapin 1 (P434L; Rao et al., 2010), showed the same results (Figure 1G-K), indicating that coalescence requires the interaction through the PRM-SH3 domain. Together, these data suggest that Dyn1xA and Syndapin 1 together form liquid-like condensates *in vitro*.

### Dyn1xA and Syndapin 1 form liquid-like condensates in COS-7 cells

Next, we tested whether Dyn1xA forms condensates with Syndapin 1 in COS-7 cells by overexpressing Dyn1xA-GFP and mCherry-Syndapin 1 (Figure 2). Similarly to the *in vitro* experiments (Figure 1), Dyn1xA and Syndapin 1 also formed spherical liquid-like condensates (Figure 2A) that underwent fusion and re-rounding upon fusion (Figure 2B). These condensates became progressively larger at 48 hours compared to 24 hours post-transfection (Figure 2A), suggesting that the expression level of proteins determines the size of the condensates. As *in vitro*, they also exhibited rapid rearrangement of molecules within condensates upon photobleaching, with their kinetics dependent on the radius of photobleaching spots (Figure 2C-E). When entire condensates were photobleached, the recovery reached ∼60% (Figure 2K), indicating that molecules can be exchanged between cytosol and puncta. Condensate formation required a specific Dyn1xA–Syndapin1 interaction, not synthetic interaction through fluorescent protein tags or other proteins natively expressed in COS-7 cells, since disrupting their interaction or Dyn1xA multimerization prevented condensates from forming (Figure S2A). Unlike with *in vitro* experiments, Syndapin 1 alone did not form condensates (Figure S2B), suggesting that endogenous proteins may interact with Syndapin 1 and prevent condensate formation. Interestingly, the expression of Dyn1xA alone led to the formation of droplet-like structures (Figure S2B). However, the kinetics of FRAP on these structures were similar between two different photobleaching radii (Figure S2C-E, 700 nm; 34.7 s, 350 nm; 30.1 s). In addition, when the entire puncta were photo-bleached, fluorescence did not recover (17.0 %, Figure S2F, G, H), suggesting that Dyn1xA alone forms aggregates in COS-7 cells. Likewise, when Dyn1xA-S774/778D and Syndapin 1 were co-expressed, the resulting condensates appeared irregular in shape (Figure 2F) and exhibited nearly identical recovery kinetics to photobleaching of two different radii (Figure 2G-I; 700 nm; 12.6 s, 350 nm; 14.9 s) and minimal recovery from full bleaching (Figure 2J-L, Dyn1xA; 60.5%, Dyn1xA-S774/778D; 18.3%). Similarly, aggregate-like puncta formed when Syndapin 1 was expressed with Dyn1xB (Figure S2I-O), which does not participate in ultrafast endocytosis or localize to the endocytic zone in neurons (Imoto et al., 2021). These results indicate that the Dyn1xA–Syndapin 1 interaction is essential for liquid-like condensate formation in COS-7 cells.

**Figure 2.**
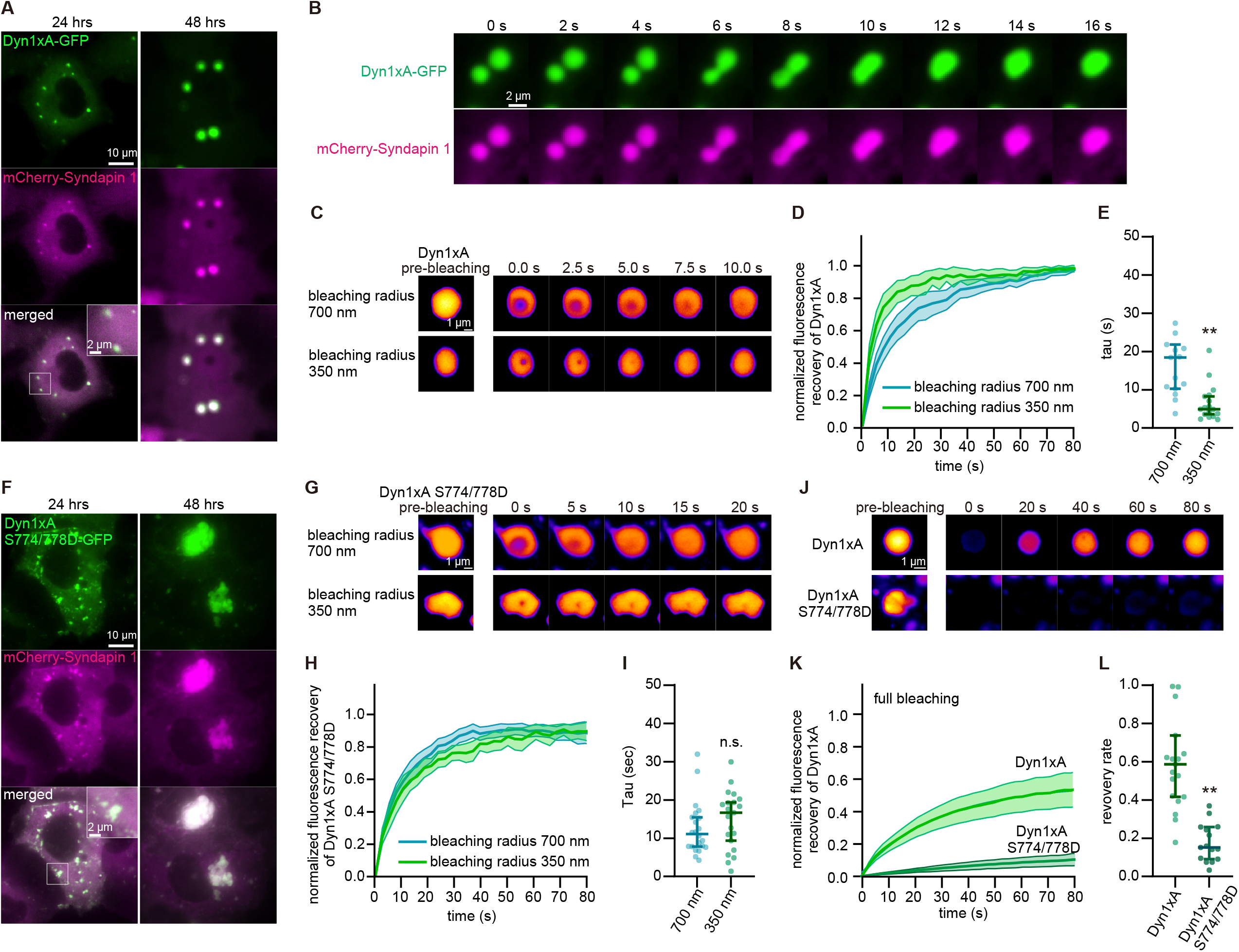
Dyn1xA exhibits liquid-like behaviors in COS-7 cells. (A) Example images of COS-7 cells expressing Dyn1xA-GFP and mCherry-Syndapin1 at 24 hours and 48 hours after the transfection. (B) Example time-lapse images showing fusion of Dyn1xA-GFP and mCherry-Syndapin1 droplets. (C) Examples time-lapse images of FRAP experiments on Dyn1xA-GFP and mCherry-Syndapin1 droplets. Time indicates after the photobleaching. Dyn1xA signals were photobleached at 480 nm using the ROI radius of 700 nm or 350 nm ROI. (D) Normalized fluorescence recovery of Dyn1xA signals in different ROI sizes. Fluorescence signals were normalized between immediately (0 s) and 80 s after the photobleaching. Times indicate after the photobleaching. The median and 95% confidential interval are shown. (E) The recovery time constant of Dyn1xA signals following the photobleaching using different ROI sizes in (D). The median and 95% confidence interval are shown. Each dot represents a Dyn1xA-Syndapin 1 droplet. p < 0.0001. Mann-Whitney test. (F) Example images of COS-7 cells expressing Dyn1xA S774/778D-GFP and mCherry-Syndapin1 at 24 hours and 48 hours after the transfection. (G) Examples time-lapse images of FRAP experiments on Dyn1xA S774/778D-GFP and mCherry-Syndapin1 droplets. Time indicates after the photobleaching. Dyn1xA signals were photobleached at 480 nm using the ROI radius of 700 nm or 350 nm. (H) Normalized fluorescence recovery of Dyn1xA signals in different ROI sizes. Fluorescence signals were normalized between immediately (0 s) and 80 s after the photobleaching. Times indicate after the photobleaching. The median and 95% confidential interval are shown. (I) The recovery time constant of Dyn1xA signals following the photobleaching using different ROI sizes in (H). The median and 95% confidence interval are shown. Each dot represents a Dyn1xA-Syndapin 1 droplet. n.s., no significance. Mann-Whitney test. (J) Examples time-lapse images of FRAP experiments on Dyn1xA and mCherry-Syndapin1 droplets or Dyn1xA S774/778D-GFP and mCherry-Syndapin1 droplets with the photobleaching laser on the entire droplets. Time indicates after the photobleaching. Dyn1xA or Dyn1xA S774/778D-GFP signals were photobleached at 480 nm. (K) Normalized fluorescence recovery of Dyn1xA or Dyn1xA S774/778D-GFP signals. Fluorescence signals were normalized between immediately (0 s) and 80 s after the photobleaching. Times indicate after the photobleaching. The median and 95% confidential interval are shown. (L) The recovery rate of Dyn1xA or Dyn1xA S774/778D-GFP signals at 80 s after the photobleaching in (K). The median and 95% confidence interval are shown. Each dot represents a Dyn1xA-Syndapin 1 droplet. p < 0.0001. Mann-Whitney test. All data are examined by n > 15 droplets from two independent cultures.

### Dyn1xA puncta display liquid-like properties in neurons

Dyn1xA forms liquid-like condensates with Syndapin 1 both *in vitro* and in COS-7 cells. To test if Dyn1xA puncta at synapses (Imoto et al., 2021) exhibit the same behavior, we first applied aliphatic alcohols, 1,6-hexanediol, 2,5-hexanediol, and 1,4-butanediol, which disrupt weak hydrophobic interactions and dissolve liquid droplets to varying degrees (Gopal et al., 2017; Kroschwald et al., 2015; Patel et al., 2007; Kozak and Kaksonen, 2019; Wilfling et al., 2020). These treatments dispersed the puncta, and Dyn1xA became diffuse along the axon within 30-60 s (Figure 3A-C). In control neurons, the puncta were stable over 1 min, as expected for the liquid-like condensates. These results are all consistent with the characteristic features of phase separation (Park et al., 2021).

**Figure 3.**
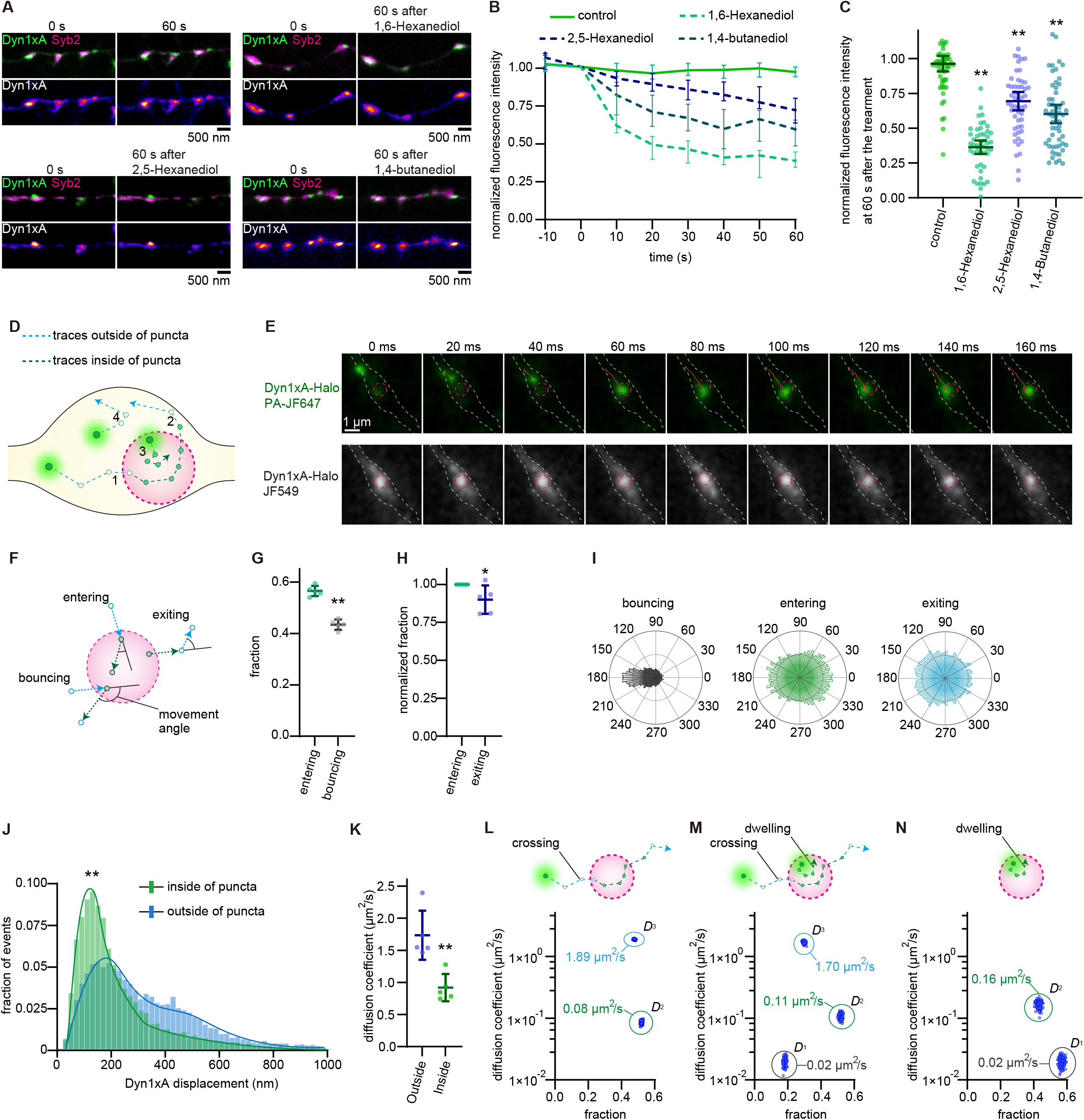
Dyn1xA exhibits liquid-like behaviors in presynapses. (A) Example time-lapse fluorescence micrographs showing Dyn1xA signals over 1 min (control), before and 1 min after the addition of 4% 1,6-hexanediol. (B) Averaged normalized fluorescence intensities of Dyn1xA puncta in control and 4% 1,6-hexanediol treatment. Fluorescence is normalized to the pre-treatment (0 s). Time indicates after the addition. The median and 95% confidence interval are shown. (C) Normalized fluorescence recovery at 60 s after treatment show in (B). p <0.0001. Mann-Whitney test. Ordinary one-way ANOVA with full pairwise comparisons by Holm-Šídák’s multiple comparisons test. The mean and SEM are shown. n > 40 presynapses for the control, the 4% 1,6-hexanediol, 4% 2,5-hexanediol or 4% 1,4-buntanediol treatment. > 3 different neurons were examined in each condition from 2 independent cultures. (D) Schematic showing hypothetical movements of Dyn1xA single molecules and their trajectories. Dyn1xA-PA-JF647 molecules are photo-activated and tracked within presynaptic boutons. The boundary of Dyn1xA puncta are visualized by Dyn1xA-JF549 (magenta dotted line). Each trajectory of Dyn1xA-PA-JF647 is generated based on its relative positions in two consecutive frames (an open circle to another). The length of each trajectory is defined as the displacement distance. All trajectories that initiate and terminate within the punctum are quantified as movement within the punctum (green dotted line). To reduce the bias imposed by the differences in available spaces between the cytosol and puncta, all trajectories outside the puncta that never cross the border of Dyn1xA puncta are not considered (light gray dotted lines) unless their next trajectory ends up in the punctum. In addition, when a trajectory crosses the boundary from outside, they are counted as the movement outside the punctum (light blue dotted line), while a trajectory crossing the boundary from inside are quantified as movement inside the punctum (green dotted line). See Materials and Methods. (E) Example time-lapse images of a single molecule of Dyn1xA in a presynaptic bouton after photo-activation. This molecule enters the Dyn1xA punctum and stays within the punctum. (F) Schematic showing hypothetical movements of Dyn1xA single molecules when they across boundary. Entering; consecutive two trajectories which moved from outside, inside, and inside. Bouncing; consecutive two trajectories which moved from outside, inside and went to outside. Exiting; consecutive two trajectories which moved from inside, outside and outside. Movement angles were calculated from the angle made by two consecutive trajectories. (G) Fraction of number of entering and bouncing events. Goes in; entering events and bouncing; 4460 events from 88 boutons, more than 10 neurons. n=5 independent cultures. p <0.0001. Mann-Whitney test. (H) Number of entering and exiting events normalized to goes in. Goes in; 5735 events [same as in (G)] and comes back; 5517 events n=5 independent cultures. p <0.0001. Mann-Whitney test. (I) Angular distribution histograms extracted from (J) and (K). (J) Histograms showing displacement distance of trajectories inside (green) or outside the puncta (light blue). (K) Transition of diffusion coefficient from outside to inside. (L) Schematic showing traces of boundary crossing and staying inside (dwelling) which were used for calculation of diffusion coefficient. Dot plot shows three components of diffusion coefficient of dyn1xA molecules (D_1_, D_2_, D_3_) detected in bouton with preaccumulations. n > 150 iterations. (M) Schematic showing a trace of boundary crossing which was used for calculation of diffusion coefficient. Dot plot shows two components of diffusion coefficient of dyn1xA molecules (D_2_, D_3_) detected in bouton with preaccumulations. n > 150 iterations. n = 5 independent cultures. 14178 and 13520 trajectories from 88 boutons and more than 10 neurons were used. p <0.0001. Mann-Whitney test.

### Dyn1xA puncta at synapses are *bona fide* liquid condensates

Although the results so far point to the liquid-like properties of the Dyn1xA puncta at synapses, they can also be explained by molecular scaffolding. In fact, both dynamin and syndapin form rigid scaffolds around endocytic vesicles during clathrin-mediated endocytosis, and they also display moderate fluorescence recovery after photo-bleaching (Taylor et al., 2012). Thus, to distinguish liquid condensates from scaffolding, we need to test whether the diffusion coefficient of molecules changes when they are inside the puncta. Molecules diffuse slower inside liquid-like condensates due to molecular crowding and intermolecular interactions (Ladouceur et al., 2020; McSwiggen et al., 2019b; Miné-Hattab et al., 2021). To this end, we performed stroboscopic photo-activatable single particle tracking (spaSPT) to monitor the behavior of individual Dyn1xA molecules within and around the puncta (McSwiggen et al., 2019b; Miné-Hattab et al., 2021).

We labeled exogenously-expressed Dyn1xA::HaloTag with JFX_549_ and PA-JF_646_ (Grimm et al., 2016, 2021) with the ratio of 2 JFX_549_:1 PA-JF_646_. With this scheme, the majority of Dyn1xA molecules are labelled with JFX_549_, visualizing the Dyn1xA puncta (Figure 3G, H, S3A, Supplementary Movie 1-3). Stochastic photo-activation of PA-JF_646_ is then used to probe the behavior of individual molecules relative to the puncta (Figure 1G, H; McSwiggen et al., 2019b; Miné-Hattab et al., 2021). Images are acquired at 50 Hz on a custom-built lattice-light sheet microscope (see STAR Methods). Custom-written MATLAB codes delineate the boundary of Dyn1xA-JFX_549_ puncta and generate trajectories of Dyn1xA-PA-JF_646_ molecules (Figure 3G; see STAR Methods for more details). The photo-activated Dyn1xA-PA-JF_646_ molecules can enter and exit the puncta, suggesting that molecules are exchanged between the cytosol and puncta. As expected for the liquid condensates (McSwiggen et al., 2019b), the molecules are trapped in the puncta upon entry while some escaping from the puncta (Figure 3D, E). Upon entry, they diffused in all directions (Figure 3F-I). Occasionally, molecules entered the puncta but bounced right back to where they came from (Figure 3F, G). Nonetheless, when they successfully entered and stayed within the puncta, their displacement distance, that is the distance traveled between two consecutive camera frames, became shorter and remained so until they exited the puncta (Figure 3J), indicating an apparent change in the diffusion coefficient. In fact, the weighted diffusion coefficient became slower after the entry into the puncta, when we fitted the histogram of the displacement distance using maximum likelihood estimation (Figure 3K). These numbers are consistent with the liquid like behavior in previous studies (Ladouceur et al., 2020; Miné-Hattab et al., 2021).

To more accurately calculate the diffusion coefficients of molecules, we used the Single-Molecule Analysis by Unsupervised Gibbs sampling (SMAUG) algorithm, which provides the number of diffusion components, their average diffusion coefficient, and the fraction of single molecules in each component (Karslake et al., 2021). The trajectories that crossed the boundary at least once (crossing) (Figure S3B, S3C) had two diffusional states: D2 (intermediate, 0.0848 ± 0.0543 µm^2^/s, 52.15 ± 0.47%) and D3 (free diffusion, 1.885 ± 0.0416 µm^2^/s, 47.85 ± 0.47 %) (Figure 3L, S3D, E), suggesting that the molecules indeed slow down when they enter the puncta. The intermediate component is consistent with the diffusion within the liquid-like environment. Interestingly, when the trajectories that initiated and terminated inside the puncta (dwelling) (Figure S3C, S3D) were included in the analysis (crossing + dwelling), three diffusional states were present (Figure 3M, S3D, S3E): D1 (slow, 0.0201 ± 0.0027 µm^2^/s, 18.24 ± 0.66 %), D2 (intermediate, 0.1066 ± 0.0103 µm^2^/s, 51.96 ± 0.72%) and D3 (free diffusion, 1.6960 ± 0.0758 µm^2^/s, 29.79 ± 0.62 %). This result suggests that some of the dwelling molecules (∼18%) are likely bound to other proteins or lipids. In fact, the analysis with only dwelling molecules showed two diffusional states: D1 (slow, 0.0197 ± 0.0002 µm^2^/s, 56.91 ± 0.09%) and D2 (intermediate, 0.1663 ± 0.0017 µm^2^/s, 43.09 ± 0.09 %) (Figure 3N, S3D, S3E), suggesting that the molecules are either bound or moving slowly within the puncta. These data indicate that the Dyn1xA puncta are liquid-like with a small fraction (∼18 %) of Dyn1xA molecules possibly bound to proteins or membranes within the puncta.

### Phase separation of Dyn1xA at synapses requires dephosphorylation of the PRM

The phosphorylation state of Dyn1xA modulates its ability to interact with Syndapin 1. Thus, altering the phosphorylation level of its PRM may affect condensate formation. To examine the requirement for dephosphorylation, we localized wild-type, phosphomimetic (S774/778D), and phospho-deficient (S774/778A) forms of Dyn1xA in neurons. We visualized Dyn1xA-GFP along with an active zone protein, Bassoon, using Stimulated Emission Depletion (STED) microscopy (Figure 2K, L). Dyn1xA signals were detected near the edge of Bassoon signals (Figure 4A). Using custom-written MATLAB codes, we defined the boundary of active zones based on Bassoon signals and quantified the distribution of Dyn1xA signals from the active zone edge (Figure 4B, See Method). Consistent with previous study (Imoto et al., 2021), most Dyn1xA signals were found within 100 nm from the active zone edge where ultrafast endocytosis takes place (Watanabe et al., 2013a). Likewise, phosphodeficient Dyn1xA accumulated within 100 nm from the active zone edge. By contrast, the phosphomimetic form was distributed broadly within boutons. We occasionally observed puncta like structure of Dyn1xA phosphomimetic form, however, FRAP experiment showed these were protein aggregate (Figure S4A, B). Since wild type Dyn1xA puncta showed FRAP recovery (Imoto et al., 2021), proper formation of Dyn1xA puncta requires hydrophobic interaction between Dyn1xA and Syndapin 1. Together, these results suggest that Dyn1xA forms liquid condensates at the endocytic zone in presynaptic boutons, and this formation is promoted by dephosphorylation.

**Figure 4.**
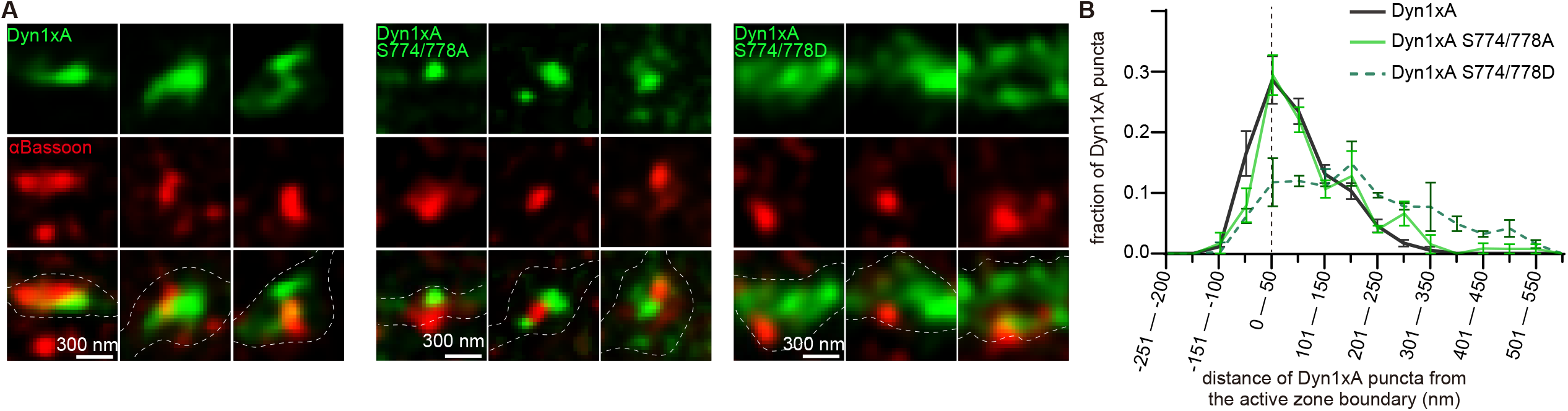
Dyn1xA condensates requires dephosphorylation of the proline-rich motif. (A) Example STED micrographs showing the localization of Dyn1xA or Dyn1xB (stained with anti-GFP antibodies) and αBassoon (stained with anti-Bassoon antibodies). (B) The distribution of Dyn1xA or Dyn1xB relative to the active zone edge. The active zone edge was defined by 50% local maxima of αBassoon signals. The median and 95% confidence interval are shown. n >25 presynapses in at least 4 neurons from two independent cultures. See Quantification and Statistical Analysis for the n values and detailed numbers.

## Discussion

### Dyn1xA liquid condensates at the endocytic zone

The majority of dynamin molecules are at first cytoplasmic and then recruited to the plasma membrane after endocytic pits are formed (Ehrlich et al., 2004; Ferguson and De Camilli, 2012; McMahon and Boucrot, 2011; Taylor et al., 2011, 2012). This recruitment depends on the SH3-containing BAR domain proteins like Syndapin 1. Once recruited, Dynamin and Syndapin 1 typically form a rigid scaffolding structure around the highly curved membrane at the neck of endocytic pits (Daumke et al., 2014; Haucke et al., 2011; Rao et al., 2010). In contrast, our recent study suggests that Dyn1xA and Syndapin 1 can also accumulate on the flat membrane prior to endocytosis (Imoto et al., 2021). In addition, molecular genetic disruption of the pre-recruited puncta formation slows down the endocytic kinetics by 100-fold. The present study now demonstrates that the pre-recruited Dyn1xA forms liquid-like condensates with Syndapin 1. Likewise, early endocytic proteins like Eps15 and Epsin also form molecular condensates to concentrate proteins at the endocytic sites for clathrin-mediated endocytosis (Day et al., 2021; Kozak and Kaksonen, 2019). Thus, molecular condensation may be a general mechanism to accumulate proteins for endocytosis.

Dyn1xA condensates may also contain an immobile fraction of molecules that are bound to either lipids or other proteins. These molecules tend to stay within the condensates and have a slower diffusion coefficient (0.0201 µm^2^/s) than the mobile fraction of molecules (0.1066 ± 0.0103 µm^2^/s). There are three possible interpretations of these immobile molecules. The first and simplest interpretation is that a certain fraction of Dyn1xA is bound to Syndapin 1 more tightly and is immobile on the plasma membrane. Consistent with this, the affinity of the Dyn1xA PRM and the Syndapin 1 SH3 is higher than a typical PRM – SH3 interaction due to their interaction at the extra residues on either side of the binding domain on Dyn1xA (Anggono and Robinson, 2007). Since the membrane interaction of Syndapin 1 is essential for their colocalization (Imoto, et al., 2021), it is quite likely that some of these proteins are more tightly bound to the plasma membrane. The second possibility is that the immobile fraction represents pre-formed rigid contractile oligomers of dynamin. In fact, Dynamin 1 can be localized to endocytic sites through its association with the clathrin lattice on flat membrane, thereby accelerating clathrin-mediated endocytosis (Srinivasan et al., 2018). However, the immobile fraction of Dyn1xA at synapses is unlikely to be analogous, since clathrin is dispensable for ultrafast endocytosis. The third possibility is that the Dyn1xA immobile fraction represents the seed of liquid-like condensates. A two-state behavior (immobile and liquid) has been observed in other liquid-like condensates. For example, stress granules have a semisolid gel-like core, which contains both mobile and immobile fractions (Jain et al., 2016; Niewidok et al., 2018). The assembly of the gel-like cores is thought to be the initial step in phase separation (Wheeler et al., 2016). Thus, immobile Dyn1xA molecules may also act as seed. Further experiments are necessary to tease apart these models.

### Liquid-like condensates as the organizing principle of synapses

Liquid-like molecular condensates concentrate and compartmentalize specific proteins and cytoplasmic materials without boundaries enforced by membranes (Banani et al., 2017; Brangwynne et al., 2009). Recent studies and our data suggest that at least three distinct condensates exist at presynapses. First, synaptic vesicles are clustered and confined within presynaptic terminals through phase separation of a peripheral synaptic vesicle protein, synapsin 1 (Milovanovic et al., 2018; Pechstein et al., 2020). Second, active zones are phase-separated through interactions between Rab3-interacting molecule (RIM) and RIM-binding protein (Wu et al., 2019). Third, our data demonstrate that Dyn1xA and syndapin 1 form molecular condensates at putative endocytic sites to drive ultrafast endocytosis. Thus, although constrained in space, presynaptic terminals are divided into three distinct domains by liquid-liquid phase separation.

This organization of multiple liquid condensates within a single compartment is similar to the recently discovered chromatin organization within the nucleus, where transcriptionally-active or inactive sites appear as distinct condensates and adhere to each other but do not coalesce (Gibson et al., 2019). Like with chromatin, liquid condensates at synapses likely maintain functionally distinct domains within a synaptic terminal as separate entities, perhaps to maintain efficient synaptic transmission. Following exocytosis, pre-recruited dynamin and other endocytic proteins can rapidly internalize the excess membranes and regenerate synaptic vesicles through ultrafast endocytosis (Watanabe et al., 2013a, 2018). The capacity of endocytosis can be tuned by simply altering the number of Dyn1xA puncta within synapses through the activity of kinases (GSK3ß, CDK-5, and Dyrk1A) and a phosphatase, calcineurin (Chen-Hwang et al., 2002; Clayton et al., 2010; Tan et al., 2003; Xue et al., 2011). Thus, these seemingly complicated events can be streamlined through the formation of functionally distinct molecular condensates, and these domains can be controlled independently through different sets of kinases and phosphatases or interdependently via calcium signaling to meet the demands of synaptic activity.

## Acknowledgements

We thank Pietro De Camilli for sharing mice and antibodies, Ira Milosevic for advice on antibodies, Geraldine Seydoux and Philip Robinson for discussion. We are also indebted to M. Delanoy, B. Smith and Hoku West-Foyle at the Johns Hopkins Microscopy Facility for technical assistance in electron and optical microscopy, Tyler Ogunmowo, Quan Gan and Sydney Brown for animal husbandry and the preparation of cells, Yuta Nihongaki for the helpful discussion on *in vitro* protein assay, Pascal Fenske, Lauren Mamer, Fereshteh Zarebidaki, Berit Söhl-Kielczynski, and Thorsten Trimbuch for assist in high-pressure freezing experiments, and Grant F. Kusick for editing of the manuscript. J. L. was supported by National Science Foundation (2105837), Johns Hopkins Catalyst award, and start-up funds from Johns Hopkins University School of Medicine. S.W. and this work were supported by start-up funds from the Johns Hopkins University School of Medicine, Johns Hopkins Discovery funds, Johns Hopkins Catalyst award, the National Science Foundation (1727260), and the National Institutes of Health (1DP2 NS111133-01 and 1R01 NS105810-01A1) awarded to S.W.. S.W. is an Alfred P. Sloan fellow, a McKnight Foundation Scholar and a Klingenstein and Simons Foundation scholar. Y.I. was supported by JSPS. M.A.C. is supported by The Wellcome Trust (204954/Z/16/Z). T.H. is an investigator of the Howard Hughes Medical Institute. T.I. and J.L. are supported by National Science Foundation (NSF 2148534).

## Author Contributions

Y.I., Y.M., and S.W. conceived the study and designed the experiments. S.W. oversaw the overall research. Y.I. performed the freezing experiments. Y.I., Y.M., and K.I. collected and analyzed the data. Y.M. wrote the analysis codes for STED and LLSM images. Y.M. and B.W. built LLSM. E.B. and M.A.C. performed the pull-down assay. Y.I., and H.M., prepared proteins for *in vitro* experiments. Y.I. and K.I. prepared neuron cultures. Y.I., K.I., H.M. generated DNA constructs for the study. J. L. carried out the theoretical estimation. Y.I and S.W. wrote the manuscript. All authors contributed to editing of the manuscript. J.L., B.W., T.H., T.I., and S.W. funded the research.

## STAR Methods

### EXPERIMENTAL MODEL AND SUBJECT DETAILS

All experiments were performed according to the rules and regulations of the National Institute of Health, USA, and the UK Animal (Scientific Procedures) Act 1986, under Project and Personal License authority (Home Office project license – 7008878). Animal protocols were approved by committee of animal care, use of the Johns Hopkins University and the University of Edinburgh Animal Welfare and Ethical Review Body. Primary cultures of mouse hippocampal neurons with the following genotypes were used in this study: C57/BL6-N; C57/BL6-J, *DNM3* KO (Raimondi et al., 2011), and *DNM1,3* DKO (Raimondi et al., 2011). The animals of the genotype *DNM1*^*+/+*^, *DNM3*^-/-^ and *DNM1*^*-/-*^, *DNM3*^-/-^ were generated by crossing *DNM1*^+/-^, *-3*^-/-^ animals. *DNM1*^*+/-*^, *DNM3*^-/-^ animals were not included in the study to avoid any confound results due to the haploinsufficiency.

### Primary neuronal cultures

To prepare primary neuronal cultures, the following procedures (Itoh et al., 2019) were carried out. Newborn or embryonic day 18 (E18) mice of both genders were decapitated. The brain is dissected from these animals and placed on ice cold dissection medium (1 x HBSS, 1 mM sodium pyruvate, 10 mM HEPES, 30 mM glucose, and 1% penicillin-streptomycin).

For high pressure freezing, hippocampal neurons were cultured on a feeder layer of astrocytes. Astrocytes were harvested from cortices with treatment of trypsin (0.05%) for 20 min at 37 °C, followed by trituration and seeding on T-75 flasks containing DMEM supplemented with 10% FBS and 0.2% penicillin-streptomycin. After 2 weeks, astrocytes were plated onto 6-mm sapphire disks (Technotrade Inc) coated with poly-D-lysine (1 mg/ml), collagen (0.6 mg/ml) and 17 mM acetic acid at a density of 13 × 10^3^ cells/cm^2^. After 1 week, astrocytes were incubated with 5-Fluoro-2′-deoxyuridine (81 µM) and uridine (204 µM) for at least 2 hours to stop the cell growth, and then medium was switched to Neurobasal-A (Gibco) supplemented with 2 mM GlutaMax, 2% B27 and 0.2% penicillin-streptomycin prior to addition of hippocampal neurons. Hippocampi were dissected under a binocular microscope and digested with papain (0.5 mg/ml) and DNase (0.01%) in the dissection medium for 25 min at 37 °C. After trituration, neurons were seeded onto astrocyte feeder layers at density of 20 × 10^3^ cells/cm^2^. Cultures were incubated at 37 °C in humidified 5% CO_2_/95% air atmosphere. At DIV14-15, neurons were used for high pressure freezing experiments.

For fluorescence imaging, dissociated hippocampal neurons were seeded on 18-mm or 25-mm coverslips coated with poly-L-lysine (1 mg/ml) in 0.1 M Tris-HCl (pH8.5) at a density of 25-40 × 10^3^ cells/cm^2^. Neurons were cultured in Neurobasal media (Gibco) supplemented with 2 mM GlutaMax, 2% B27, 5% horse serum and 1% penicillin-streptomycin at 37 °C in 5% CO_2_. Next day, medium was switched to Neurobasal with 2 mM GlutaMax and 2% B27 (NM0), and neurons maintained thereafter in this medium. For biochemical experiments, dissociated cortical neurons were seeded on poly-L-lysine coated plates with Neurobasal media supplemented with 2 mM GlutaMax, 2% B27, 5% horse serum and 1% penicillin-streptomycin, at a density of 1 × 10^5^ cells/cm^2^. Next day, the medium was switched to Neurobasal medium with 2 mM GlutaMax and 2% B27, and neurons maintained in this medium thereafter. A half of the medium was refreshed every week.

### Expression constructs

All the plasmids used in this study and primers to make these construct was listed in Table S1. DNA cloning was performed using transformation into competent DH5α cells. For the dynamin rescue constructs, wild-type Dyn1xA from human sequences were amplified from plasmids (generously provided by Pietro De Camilli’s lab) and fused by Gibson assembly (NEW ENGLAND BioLabs) after a self-cleaving P2A site within a pFUGW derived plasmid, which encodes nuclear RFP controlled by the human synapsin 1 promoter (f(syn)NLS-RFP-P2A), to form f(syn)NLS-RFP-P2A-Dyn1xA. Using the same approach, we generated Dyn1xA S774/778A, S774/778D, Dyn1xB S774/778A.

Dyn1xA-GFP was purchased from addgene. To generate Dyn1xA S774/778A-GFP and Dyn1xA-S774/778D GFP, Dyn1xA-GFP construct was linearized using the primers with corresponding mutations and were re-circularized using In-Fusion HD cloning kit (TAKARA). To generate Dyn1xA-GFP_mCherry-Syb2, rat Syb2 (addgene) and mCherry (addgene) sequence were amplified and combined with linearized Dyn1xA construct using In-Fusion HD cloning kit. As a result, mCherry-Syb2 sequence are inserted in to downstream of Dyn1xA-GFP sequence. To generate cytosolic tdTomato, tdTomato sequence was amplified from tdTomato vector (Clontech) and was ligated at the XhoI and KpnI digestion sites of pCAGGS construct by DNA ligation kit Solution I (TAKARA).

For pull-down assays, wild-type rat Dyn1xA-PRM (C-terminus residues 746-864), S774/778E and S774/778A mutants were generated by amplifying the required region from Dyn1aa-GFP (rat) in pEGFP-N1. The amplified product was inserted into pGEX4T-1 vector (Amersham Biosciences) using the restriction enzymes EcoRI and NotI (underlined in sequence). Subsequent modifications to generate S774/778E and S774/778A mutants used the QuickChange site-directed mutagenesis kit (Stratagene) and were confirmed by DNA sequencing (Anggono et al., 2006). Wild-type rat Dynamin-1xb C-terminus (residues 746-851) was generated by amplifying the required region from Dyn1ab in a pCR3.1 expression vector (Cao et al 1998). Similar to Dyn1xA, the amplified product was inserted into a pGEX-4T-1 using the restriction enzymes EcoRI and NotI (Xue et al., 2011).

For the recombinant protein expression constructs, wile-type Syndapin 1 and Syndapin 1 P434L sequences were amplified from mCherry-shRNA resistant Syndapin 1 and mCherry-shRNA resistant Syndapin 1 P434L, respectively. In this design, the forward primer inserts a BamHI site and TEV cleavage sequence (ENLYFQ/SG) in the upstream of Syndapin 1, and the reverse primer inserts a NotI site after the stop codon of Syndapin 1. The PCR products and pET28a were digested by BamHI-HF and NotI-HF, and ligated by T4 ligase (NEB, #M0202). Wild-type Dyn1xA and Dyn1xA S774/778D sequences were amplified from Dyn1xA -GFP and Dyn1xA S774/778A-GFP, respectively. Then, combined with linearized pGex6T using In-Fusion HD cloning kit.

### Lentivirus production and infection

For lentivirus production, HEK293T cells were maintained in DMEM supplemented with 10% FBS and 0.2% penicillin-streptomycin. One day after plating 6.5 × 10^6^ cells in T75 flask (Corning), medium was replaced with Neurobasal-A media supplemented with 2 mM GlutaMax, 2% B27 and 0.2% penicillin-streptomycin. Cells were transfected using polyethylenimine with a pFUGW plasmid encoding the RFP co-infection marker and insert of interest, and two helper plasmids (pHR-CMV8.2 deltaR and pCMV-VSVG) at a 4:3:2 molar ratio. Three days after transfection, supernatant was collected and concentrated to 20-fold using Amicon Ultra-15 10K centrifuge filter (Millipore). ChetaTC and dynamin rescue viruses were added to each well of neurons at DIV 3 with 15 µl per well (12-well plates). Syndapin 1 shRNA virus was added to each well of neurons at DIV 3 with 5 µl per well (12-well plates, 75 K neurons / well). Syndapin 1 rescue viruses were added to each well of neurons at DIV 3 with 5 µl per well (12-well plates, 75K neurons / well) or 20 µl per well (6-well plates, 300 K neurons / well). In all cases, infection efficiency of over 95% was achieved. Neurons were fixed or used for high pressure freezing at DIV13-16.

### Transient transfection of neurons

For transient protein expression, neurons were transfected at DIV13-14 using Lipofectamine 2000 (Invitrogen) in accordance with manufacture’s manual with minor modifications (Araki et al., 2015). Before transfections, a half of medium in each well was transferred to 15 mL tubes and mixed with the same volume of fresh NM0 and warmed to 37 °C in CO_2_ incubator. This solution later served as a conditioned medium. Briefly, 0.25-2 µg of plasmids was mixed well with 2 µl Lipofectamine in 100 µl Neurobasal media and incubated for 20 min. For Dyn1 expression, 0.5 µg of constructs were used to reduce the expression level. For tdTomato expression, 2.0 µg of constructs were used. For Syndapin 1 expression, 0.25 µg of constructs were used in the Syndapin 1 knock-down neurons. The plasmid mixture was added to each well with 1 ml of fresh Neurobasal media supplemented with 2 mM GlutaMax and 2% B27. After 4 hours, medium was replaced with the pre-warmed conditioned media. Neurons were incubated for less than 20 hours and fixed for imaging or subjected to live imaging. For GFP knock-in experiments, hippocampal neurons were transfected at DIV 3 with pORANGE construct (1.5 µg) and pmCherry-N1 (0.5 µg) as cell-fill using Lipofectamine 2000 and fixed at DIV15-18.

### Immunofluorescence staining

For immunofluorescence, 125 k neurons were seeded on 18 mm poly-L-lysins (1 mg/mL, overnight) coated coverslips (Thickness 0.09-0.12 mm, Caroline Biological) in 12-well plate (Corning). Neurons were fixed at DIV14-15 with pre-warmed (37 ºC) 4% paraformaldehyde and 4% sucrose in PBS for 20 min and permeabilized with 0.2% Triton X-100 in PBS for 8 min at room temperature. After blocking with 1% BSA in PBS for 30 min, cells were incubated with primary antibodies diluted in 1% BSA/PBS overnight at 4 °C, followed by appropriate secondary antibodies diluted 1:400 in 1% BSA/PBS for 1 hour at room temperature. Coverslips were rinsed with PBS three times and were mounted in ProLong Gold Antifade Mountant (Invitrogen) and stored at 4 ºC until imaging. For the STED imaging, Dyn1xA-GFP or Dyn1xB-GFP was stained with 1:500 dilution of anti-GFP antibody, Rabbit polyclonal (MBL International) and endogenous Bassoon was stained with 1:500 dilution of anti-Bassoon antibody, mouse monoclonal (Synaptic Systems). 50 µM of anti-rabbit ATTO647 (Rockland) and anti-mouse Alexa594 (Invitrogen) was used for the secondary antibodies. Coverslips were rinsed with PBS three times and mounted in ProLong Diamond Antifade Mountant (Thermo Fisher) and stored at 4 ºC until imaging.

### Confocal microscopy imaging and analysis

For fluorescence imaging, all samples were imaged using a confocal microscope Zeiss LSM880 (Carl Zeiss). Fluorescence was acquired using a 63x objective lens (NA = 1.4) at 2048×2048 pixel resolution with the following settings: pixel Dwell 1.02 µs and pin hole size at 2 airy unit. For experiments comparing fluorescence intensity, these settings were left constant between samples. Snapshots were captured at the cell body first and then 3-7 different locations along the axon of the same neurons, typically covering 1-3 mm length from each neuron. All Dyn1-GFP transfected neurons in each coverglass were imaged. Axons were distinguished from dendritic processes based on their morphology (thin and lacking spines). If GFP signals were aberrantly saturated throughout the processes, they were dismissed to avoid overexpression artifact. Apparently dying neurons and glia cells were also excluded. We quantified all presynaptic varicosities along axons in each image. Presynaptic regions were confirmed with the Syb2-Alexa647 signals. The Syb2-Alexa647 signals were used to define the boundaries of regions-of-interest (ROIs) for quantifications, and all Dyn1-GFP and mCherry-Syndapin 1 signals within ROIs were measured as the total signals at each synapse. To define puncta within the boutons, we applied Gaussian smoothing (*σ* = 0.2) on images, performed threshold cut-off on the average fluorescence intensity of Dyn1xA and Syndapin 1 in the intersynaptic axonal regions. The circumference of each punctum adjacent to or within Syb2 signals were delineated and set as ROIs, and fluorescence intensity measured. The background signals were subtracted using the same size of ROI but 40-60 pixels away from the original ROI and outside of neural processes. For the quantification of the colocalization between Dyn1xA and Syndapin 1 puncta, ROIs of overlapped Dyn1xA and Syndapin 1 were counted. Fluorescence intensity was normalized to the average signal within the corresponding cell body of each axon. False colored images were made by applying LUT colors and ratio signal intensities are measured by calibrate bar function in Fiji.

### Live imaging and analysis

For live-cell imaging, 250 k neurons were seeded on 25 mm coverslips (thickness 0.13-0.17 mm, Carolina Biological) in 6-well plate (Corning). Dyn1xA-GFP and mCherry-Syb2 were expressed on the same construct using pDyn1xA-GFP_P2A_mCherry-Syb2 construct by transient transfection. For the digitonin treatment, 10 mg/mL of digitonin (Sigma-Aldrich) was prepared in MilliQ water and diluted into final concentration 5 µg/mL or 500 µg/mL in NM0. For the 1,6-Hexanediol treatment, 4 % of 1,6-Hexanediol was directly dissolved in NM0. For imaging, cover slips were transferred to live cell round chamber (ALA science) filled with NM0. FK506 (Tocris) or CHIR99021 (Sigma-Aldrich) were dissolved in DMSO and added to the imaging chamber to achieve the final concentrations of 2 µM and 10 µM, respectively, 30 min before imaging. 0.05 % of DMSO were added to the chamber for control experiments. All live imaging experiments were carried out on a Zeiss LSM880 confocal microscope at 37 °C in humidified 5% CO_2_/95% air atmosphere, and live imaging data was acquired by raster scan.

For photobleaching experiments, 33 frames were collected with a time interval of 2.5 s. After 3 frames, the maximum laser power was applied to bleach fluorescence. The 4th frame was started immediately after bleaching. 30 additional frames were collected after bleaching. A circular ROI was placed around each Dyn1xA punctum found adjacent to the mCherry-Syb2 signal, and Dyn1xA-GFP signals within the ROI was photobleached with a 488-nm laser. For quantifications, average fluorescence intensity was measured from each ROI and a random region outside of the cell, which was used for the background subtraction. A circular ROI of the same size was used to measure signals from an unbleached region to measure the degree of imaging-related photobleaching. The fluorescence intensity at the ROIs was normalized to the intensity before bleaching (3rd frame) and after bleaching (4th frame). To correct signal loss due to the bleaching during the imaging, the normalized fluorescence recovery over time, *F*(*t*), was then fitted to an exponential function:

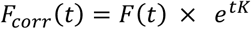

where the corrected normalized fluorescence over time is *F*_*corr*_ (*t*), time after the photo bleaching is *t*, and *K* is the rate constant acquired from the unbleached region and measured exponential curve fitting tool in Fiji.

The *F*_*corr*_ (*t*), was then fitted to an exponential function using GraphPad Prism (v8);

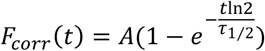

where the recovery is *A*, half-recovery time is 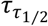.

The diffusion coefficient, D, was fitted to an,

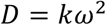

where the rate constant acquired from the bleached region is *k* and *ω* is the diameter of the bleached region.

For the cytosolic extraction experiments with digitonin, a total of 17 frames were collected with a time interval of 5 s: 5 frames were collected prior to the application of digitonin (500 µg/mL in NM0)(Sigma-Aldrich), and additional 12 frames were collected thereafter. For the 1,6,-hexanediol treatment, a total of 12 frames were collected with a time interval of 10 s: 2 frames before the addition of 4 % 1,6,-hexanediol (Sigma-Aldrich) in NM0 and 10 frames afterwards. For the purpose of imaging-related photobleaching corrections, the same number of frames were collected at additional regions distant from the imaged before the experiments. For quantifications, the average fluorescence intensities before and after the treatment were measured within the same ROIs, which were selected as described in the previous section: one ROI covering the entire Syb2 signals and another ROI along the circumference of the Dyn1xA-GFP puncta. For presentations, the brightness and the contrast in each image were adjusted in Fiji and cropped in Adobe photoshop 2021 (Adobe).

### Stimulated emission depletion microscopy (STED) imaging

All STED images were obtained using a home-built two-color STED microscope (Han and Ha, 2015). Basically, a femtosecond laser beam with repetition rate of 80 MHz from a Ti:Sapphire laser head (Mai Tai HP, Spectra-Physics) is split into two parts: one part is used for producing the excitation beam, which is coupled into a photonic crystal fiber (Newport) for wide-spectrum light generation and is further filtered by a frequency-tunable acoustic optical tunable filter (AA Opto-Electronic) for multi-color excitation. The other part of the laser pulse is temporally stretched to ∼300 ps (with two 15-cm-long glass rods and a 100-m long polarization-maintaining single-mode fiber, OZ optics), collimated and expanded, and wave-front modulated with a vortex phase plate (VPP-1, RPC photonics) for hollow STED spot generation to de-excite the fluorophores at the periphery of the excitation focus, thus improving the lateral resolution. The STED beam is set at 765 nm with power of 120 mW at back focal plane of the objective (NA=1.4 HCX PL APO 100×, Leica), and the excitation wavelengths are set as 594 nm and 650 nm for imaging Alexa-594 and Atto-647N labeled targets, respectively. The fluorescent photons are detected by two avalanche photodiodes (SPCM-AQR-14-FC, Perkin Elmer). The images are obtained by scanning a piezo-controlled stage (Max311D, Thorlabs) controlled with the Imspector data acquisition program.

### STED image analysis

A custom MATLAB code package was used to analyze the endocytic protein distribution relative to the active zone marked by Bassoon in STED images. First, the STED images were blurred with a Gaussian filter with radius of 1.2 pixels to reduce the Poisson noise, and then deconvoluted twice using the built-in deconvblind function: the first point spread function (PSF) input is measured from the unspecific antibodies in the STED images, and the second PSF input is chosen as the returned PSF from the first run of blind deconvolution (Sapoznik et al., 2020). Each time 10 iterations are performed. All of presynaptic boutons in each deconvoluted images were selected within 30×30 pixel (0.81 µm^2^) ROIs based on varicosity shape and Bassoon signals. The active zone boundary was identified as the contour that represents half of the intensity of each local intensity peak in the Bassoon channel, and the Dyn1 clusters are picked as the local maxima. The distances between the Dyn1 cluster centers and active zone boundary are automatically calculated. Dyn1 clusters over crossing the ROIs and the Bassoon signals outside of the transfected neurons were excluded from the analysis. The MATLAB scripts are available upon request.

### Lattice light sheet microscopy (LLSM) imaging

For LLMS imaging, 120 k neurons were seeded on 18 mm coverslips (thickness 0.13-0.17 mm, Carolina Biological) in 12-well plate (Corning). Dyn1xA-Halo was expressed transient transfection at DIV13. At DIV14, 100 nM JF-549 (Grimm et al., 2021) and 50 nM JF-PA-647 (Grimm et al., 2016) (Gifts from Janelia Research Campus) were incubated at 37 °C for 30 min in 1 ml NM0 followed by 4 times wash (30 min each) in the 12-well plate. Before imaging, a coverslip was rinsed three times by pre-warmed imaging buffer [140 mM NaCl, 2.4 mM KCl, 10 mM HEPES, 10 mM Glucose (pH adjusted to 7.3 with NaOH, 300 mOsm, 1 mM CaCl_2_, 4 mM MgCl_2_, 3mM NBQX and 30 mM Bicuculline]. All these processes were performed under dark room to reduce pre-activated JF-PA647. For the LLSM imaging, a home-built LLSM setup was used (Chen et al., 2014). The detailed blueprints are provided by the Betzig group at the Howard Hughes Medical Institute Janelia Research Campus. Basically, four laser beams (405nm/250mW, RPMC; 488nm/300 mW, MPB Communications; 560nm/500 mW, MPB Communications and 642nm/500 mW, MPB Communications) were expanded and collimated independently to a size of 2.5 mm, and then were combined and sent into an acoustic optical tunable filter (AA Opto-Electronic) for rapid channel switching and power adjustment. To achieve single particle detection sensitivity, two pairs of cylindrical lenses (25mm /200 mm and 250mm/50 mm) were then used to convert the Gaussian beam into a thin stripe illumination projected onto a binary phase spatial light modulator (SLM, Forth Dimension, SXGA-3DM). The squared lattice patterns with an inner NA of 0.44 and outer NA of 0.55 was projected on the SLM, which is relayed by a 500-mm lens onto an annular mask conjugated to the back pupil plane of the excitation (Special Optics, 0.65 NA, 3.74 mm WD). Two galvo mirrors (Cambridge Technology, 6215H) were used for the excitation light sheet positioning along the axis of the detection objective (Nikon, CFI Apo LWD 25XW, 1.1 NA, 2 mm WD) and dithering along the orthogonal direction. The power of the 642-nm excitation beam was measured at the back pupil plane of the excitation objective are 3.3 mW. The emitted fluorescence was collected by the detection objective (Nikon, CFI Apo LWD 25XW, 1.1 NA, 2 mm WD) orthogonally mounted and projected onto an sCMOS camera (Hamamatsu, Orca Flash 4.0 v3) by a 500-mm tube lens. A quad-band filter (Semrock) was placed before the camera. Single-particle tracking was recorded by illuminating the sample by the 642-nm laser with 20-ms exposure time interleaved with 1-2 ms 405-nm activation for 300 frames. 10 frames with 560-nm excitation were collected before and after recording the single particle dynamics for generating the Dyn1xA pre-accumulation mask channel.

### Single particle tracking analysis

A custom MATLAB code package (available upon request) was used to analyze single particle dynamics of the Dyn1xA. Source data including all trajectory files are available from sharing server (https://data.mendeley.com/datasets/bkn78y232y/draft?preview=1). First, the individual particles were detected and linked into trajectories by running the Fiji plugin TrackMate (Tinevez et al., 2017) in batch to generate xml files recording the spatiotemporal coordinates of each particle in each trajectory. The xml files were then read by the customized MATLAB code. To generate the mask boundaries representing the pre-accumulation of Dyn1xA, the 560-nm channel images were averaged, and the boundaries were selected as the contours with 65% of the local intensity maxima. Only the trajectories that have overlap with the pre-accumulation mask were selected and further analyzed.

To calculate the weighted diffusion coefficient inside and outside the Dyn1 mask, the displacements with the start and end point purely inside and outside the mask were collected and fitted with the following distribution with 3 diffusive states using maximum likelihood estimation:

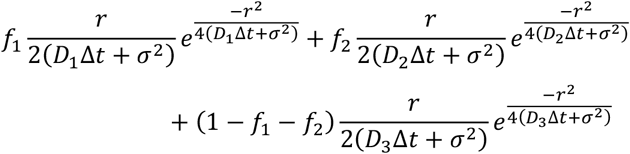

where D_1_, D_2_ and D_3_ are the diffusion coefficients with the fraction of f_1_, f_2_ and f_3_. *Δt* is the exposure time, and *σ* is the localization precision. The weighted diffusion coefficient was calculated as *f*_1_*D*_1_ + *f*_2_*D*_2_ + (1 − *f*_1_ − *f*_2_)*D*_3_

To extract the full information within single-particle trajectories collected in our experiments, which potentially contain multiple interconverting species with different diffusive state, we also applied the unsupervised Gibbs sampling (SMAUG) algorithm using nonparametric Bayesian statistics with the default settings (Karslake et al., 2021).

### Protein expression and GST-pulldown assay

Isolated nerve terminals, or synaptosomes, were prepared from rat brains of both sexes (Cousin and Robinson 2000). GST fusion proteins were expressed in bacteria at 19 °C for 16-18 hours in 500 ml cultures of supermedia (5g/L NaCl, 25g/L yeast extract (Alfa Aesar J60287), 15g/L Tryptone (Merk 1.11931), pH7.4) and coupled to glutathione-Sepharose beads (GE healthcare, 17-0756-01).

Nerve terminals were solubilized for 5 minutes at 4 °C in 25 mM Tris, pH 7.4, with 1% Triton X-100, 150 mM NaCl, 1 mM EGTA, 1 mM EDTA, 1 mM PMSF and protease inhibitor cocktail (Sigma, P58849) and centrifuged at 20,442 g for 5 minutes at 4 °C. The subsequent supernatant was incubated with GST-fusion proteins for 2 hours at 4 °C under constant rotation. After washing in lysis buffer (including a 500 mM NaCl wash), beads were washed twice in cold 20 mM Tris (pH 7.4) and boiled in SDS sample buffer (67 mM SDS, 2 mM EGTA, 9.3% glycerol, 12% β-mercaptoethanol, 0.7 mg/mL Bromophenol blue, 67 mM Tris-HCl, pH 6.8). The released proteins were separated by SDS-PAGE and analyzed by Western blotting. The blots were blocked for 1 h at room temperature in 5 % semi-skimmed milk powder dissolved in PBS-0.5 % PVP-40 before incubation in primary antibodies in PBS-0.5% PVP40 (P0930, Merck) overnight at 4 °C under constant rotation (anti-syndapin/PACSIN1, Abcam, ab137390, 1:5000; anti-GST, 27-4577-01, 1:10.000, Merck.). Blots were washed in PBS (pH 7.4) and incubated with secondary antibodies, either donkey anti-rabbit-HRP conjugated (1:20000, NBP1-7185) or donkey anti-goat-HRP (1:20000, NBP1-74815), for 2h. Enhanced Chemiluminescence substrate (Thermo Scientific, 10590624) was applied to the blots prior to exposure to medical X-ray film (Konica Minolta, MG SR, 23361). The films were developed using an Ecomax X-ray developer (S/N 118630) with band densities from scanned images quantified using Fiji (ImageJ) software. Syndapin bands were normalized to the GST fusion protein band.

### Protein expression and purification for *in vitro* phase separation assay

Expression and purification of dynamin proteins were performed as previously described with minor modifications (Imoto et al., 2018). Wild-type Dyn1xA and Dyn1xA S774/778D constructs were transformed into *E. coli* BL21-CodonPlus (DE3)-RIPL (Agilent Technologies, # 230280). Transformed BL21-CodonPlus strain cells were cultured at 37 °C for 12 hours in 200 mL of LB medium, scaled up to 2 L of LB medium, and further incubated at 37 °C for 2 hours and at 18 °C for 1 hour. IPTG was added to a final concentration of 0.1 mM, and cells were harvested after a further 12-h incubation at 18 °C by centrifugation at 5000×g for 15 min. Cell pellets were resuspended in 200 mL of HEPES buffer (HDB800) containing 800 mM NaCl, 20mM HEPES-NaOH, pH 7.5, 2 mM EGTA, 1mM MgCl_2_, 1 mM DTT, and a cOmplete protease inhibitor cocktail. Resuspended pellets were lysed by the microfluidizer (80 psi, 3 times). Soluble fraction was collected by centrifugation at 20,000× g, 4 °C, 30 min and rotated at 4 °C with 0.5 mL bed volume of glutathione Sepharose 4B beads (GE Healthcare) for 1 hour. The beads were collected by centrifugation at 1,500× g, 4 °C, 1 min and resuspended in 25 mL of HBD800. The beads were collected again by the centrifugation and resuspended in 25 mL of cOmplete-free HDB800. Then, beads resuspension was loaded onto a 10 mL Poly-Prep Chromatography column (BioRad) and washed with 25 mL of cOmplete-free HDB800 at 4 °C. After the washing, Sepharose beads were treated with 20 µL of PreScission Protease (GE Healthcare) in 5 mL of cOmplete-free HDB800 at 4 °C for 2 days and the cleaved-off proteins were eluted. Expression of wild-type and P434L Syndapin 1 was performed in *E. coli* BL21-CodonPlus (DE3)-RIPL (Agilent Technologies, # 230280). After overnight culture in LB media containing Kanamycin and Chloramphenicol (LB/Kan/Cam), 5 mL preculture was inoculated to 1 L LB/Kan/Cam and cultured at 37 °C for 4 hours. At OD600 = 0.450–0.550, 500 µM IPTG was added, and the proteins expressed at 23 °C for 26 hours.

Two-liter culture cell pellet was resuspended by 75 mL of buffer S (S for syndapin) [50 mM HEPES-NaOH (pH 7.5 at room temperature), 200 mM NaCl, 1 mM DTT] supplemented with 5 mM MgCl_2_, 1 mM PMSF, and cOmplete protease inhibitor cocktail (Roche, #4693159001) (1 tablet/75 mL) and lysed by microfluidizer (Microfluidics, Model M-110Y, 80 psi, 3 times). Soluble fraction was collected by centrifugation at 10,000× g, 4 °C, 60 min and applied to pre-equilibrated Ni-NTA agarose (QIAGEN, #30230, 3 mL resin/2 L culture). The proteins were washed and eluted by the stepwise imidazole gradient in Buffer S at 20, 30, 40, 50, 100, 200, and 500 mM Imidazole; 3 column volume each. Forty to five hundred imidazole fractions were collected and mixed with 1/50 weight of 6xHis-MBP-TEV-protease-S219V (purified in house) to remove N-terminal 6xHis tag of Syndapin 1 during 2 overnight dialysis against 25 mM HEPES-NaOH (pH 7.5 at room temperature), 200 mM NaCl, 1 mM DTT. His-tag-cleaved Synapin-1 was separated from un-cleaved sample and TEV protease by passing through Ni-NTA agarose resin (3 mL) and diluted into half with 25 mM HEPES-NaOH (pH 7.5) 1 mM DTT. The protein was further applied to pre-equilibrated HiTrapQ anion exchange column (Cytiva, #17115301, 1 mL) and eluted by linear gradient of NaCl from 100 mM to 500 mM. Higher purity fractions were then collected, concentrated by Amicon-Ultra (Millipore, #UP901024) (10 kDa cut off), snap frozen, and stored at -80 ◦C.

Proteins were concentrated by Amicon-Ultra (Millipore, #UP901024) (10 kDa cut off), snap frozen, and stored at -80 ◦C. The purity of wild-type Syndapin 1, P434L, w-type Dyn1xA and Dyn1xA S774/778D was evaluated by SDS-PAGE.

### Protein labelling

Proteins were labelled using Alexa Fluor™ 488 or Alexa Fluor™ 568 NHS Ester (Succinimidyl Ester) (Thermo Fisher) in HDB. Dye and proteins were reacted at 2;1 molar ratio. Reaction mixtures were rotated for 60 min at room temperature, then labelled protein was separated from unconjugated dye using Zeba™ Spin Desalting Columns, 7K MWCO, 0.5 mL (ThermoFisher). Reaction and the column washing buffers were as same as protein elution buffer of Dynamin 1xA or Syndapin 1.

### *In vitro* phase separation assay

In most of the experiments, Dyn1xA or Dyn1xA-S774/778D was used at 2 µM, which was the maximum concentration without having protein aggregates. Syndapin1 or Syndapin1 P434L was used at 20 µM, reflecting physiological concentration within synapses (Wilhelm et al., 2014). Before experiments, the proteins were centrifuged at 15,000× rpm at 4 °C for 10 min to remove aggregates. For the droplet formation assay, Dyn1xA and Syndapin1 are mixed at room temperature in 50 µl of HDB200 followed by addition of 5% PEG (Sigma) to their final concentrations. After 5min incubation, reaction mixture was transferred to 8-well glass chamber (LAB-TEK). The fluorescence images were collected at room temperature using 100x objective lens (NA=1.45) and LED light source at 2048×2048 pixel resolution by Nikon ECLIPSE Ti2 equipped with ORCA-Fusion BT Digital CMOS camera (HAMAMATSU). NIS-Elements AR software were used for the image acquisition. For the live imaging of the droplet fusion, confocal microscope Zeiss LSM880 was used as described above. For the sedimentation assay, 5% PEG was added to protein mixtures with the total volume of 100 µl. After the incubation at room temperature for 5 min, the protein mixtures were centrifuged 15,000× rpm for 10 min. Pellets were dissolved in 100 µl of SDS sample buffer. Supernatant and pellet fractions were examined by 4-20% SDS-PAGE gels (Biorad) and gel code staining (Thermo Fisher). For the FRAP experiments and analysis, images were acquired at room temperature using confocal microscope Zeiss LSM880 as described above. Images were acquired at 2.5 s/frame, and photobleaching with the 488 nm laser was performed just after the third frame. Droplets of similar size were selected for the experiments. For the FRAP using different ROI size, ROIs radius were setup as small as possible compared to droplet radius, which was enough to ignore influx of fluorescent molecules from outside. The intensity range was normalized to maximum and minimum intensity values after post-bleaching frames. For the bleaching entire droplet, the ROI with same radius as the droplets was used. The intensity range was normalized to the 1^st^ frame of post-bleaching images.

### Phase separation assay in COS-7 cells

COS-7 cells were seeded at 80k on 18 mm cover slips (thickness 0.13-0.17 mm, Carolina Biological) coated with 100 µg/ml fibronectin (Sigma). On next day, the 0.5 µg each of plasmid DNA were transfected using Lipofectamine (Invitrogen) in accordance with manufacture’s manual. At ∼24 or ∼48 hours post transfection, coverslips were transferred to live cell round chamber (ALA science) filled with DMEM. FRAP experiments and analysis were performed at 37 °C using a confocal microscope Zeiss LSM880 as described above.

### Flash-and-freeze experiments

For flash-and-freeze experiments (Watanabe et al., 2013b, 2014). sapphire disks with cultured neurons (DIV14-15) were mounted in the freezing chamber of the high-pressure freezer (HPM100 or EM ICE, Leica), which was set at 37 ºC. The physiological saline solution contained 140 mM NaCl, 2.4 mM KCl, 10 mM HEPES, 10 mM Glucose (pH adjusted to 7.3 with NaOH, 300 mOsm, 4 mM CaCl_2_, and 1 mM MgCl_2_. Additionally, NBQX (3 mM) and Bicuculline (30 mM) were added to suppress recurrent network activity following optogenetic stimulation of neurons. To minimize the exposure to room temperature, solutions were kept at 37 ºC water bath prior to use. The table attached to the high-pressure freezer was heated to 37 ºC while mounting specimens on the high-pressure freezer. The transparent polycarbonate sample cartridges were also warmed to 37 ºC. Immediately after the sapphire disk was mounted on the sample holder, recording solution kept at 37 ºC was applied to the specimen and the cartridge was inserted into the freezing chamber. The specimens were left in the chamber for 30 s to recover from the exposure to ambient light. We applied a single light pulse (10 ms) to the specimens (20 mW/mm^2^). This stimulation protocol was chosen based on the results from previous experiments showing approximately 90% of cells fire at least one action potential (Watanabe et al., 2013a). The non-stimulation controls for each experiment were always frozen on the same day from the same culture. We set the device such that the samples were frozen at 1 or 10 s after the initiation of the first stimulus. For ferritin-loading experiments, cationized ferritin (Sigma-Aldrich) was added in the saline solution at 0.25 mg/ml. The calcium concentration was reduced to 1mM to suppress spontaneous activity during the loading. The cells were incubated in the solution for 5 min at 37 ºC. After ferritin incubation, the cells were immersed in the saline solution containing 4 mM Ca^2+^. For dynamin experiments, all samples were incubated with 1 µM TTX for overnight to block spontaneous network activity and reduce the number of pits arrested on the plasma membrane prior to flash-and-freeze experiments (Raimondi et al., 2011; Wu et al., 2014).

Following high-pressure freezing, samples were transferred into vials containing 1% osmium tetroxide (EMS),1% glutaraldehyde (EMS), and 1% milliQ water, in anhydrous acetone (EMS). The freeze-substitution was performed in an automated freeze-substitution device (AFS2, Leica) with the following program: -90C for 5–7 h, 5C per hour to -20 ºC, 12 hours at -20 ºC, and 10 ºC per hour to 20 ºC. Following *en bloc* staining with 0.1% uranyl acetate and infiltration with plastic (30% for 3 hours, 70% for 4 hours, and 90% for overnight), the samples were embedded into 100% Epon-Araldite resin (Araldite 4.4 g, Epon 6.2 g, DDSA 12.2 g, and BDMA 0.8 ml) and cured for 48 hours in a 60 ºC oven. Serial 40-nm sections were cut using a microtome (Leica) and collected onto pioloform-coated single-slot grids. Sections were stained with 2.5% uranyl acetate before imaging.

### Electron microscopy

Ultrathin sections of samples were imaged at 80 kV on the Philips CM120 at 93,000x magnification or on the Hitachi 7600 at 150,000x. At these magnifications, one synapse essentially occupies the whole screen, and thus, with the bidirectional raster scanning of a section, it is difficult to pick certain synapses, reducing bias while collecting the data. In all cases, microscopists were additionally blinded to specimens and conditions of experiments. Both microscopes were equipped with an AMT XR80 camera, which is operated by AMT capture software v6. About 120 images per sample were acquired. If synapses do not contain a prominent postsynaptic density, they were excluded from the analysis – typically a few images from each sample fall into this category.

### Analysis of electron micrographs

Electron micrographs were analyzed using SynpasEM. Briefly, images were pooled into a single folder from one set of experiments, randomized, and annotated blind. Using custom macros (Watanabe et al., 2020), the x- and y-coordinates of the following features were recorded in Fiji and exported as text files: plasma membrane, postsynaptic density, synaptic vesicles, large vesicles, endosomes, and pits. By visual inspection, large vesicles were defined as any vesicle with a diameter of 60-100 nm. Endosomes were distinguished by any circular structures larger than large vesicles or irregular membrane-bound structures that were not mitochondria or endoplamic reticulum. Late endosomes and multivesicular bodies were not annotated in this study. Pits were defined as smooth membrane invaginations within or next to the active zone, which were not mirrored by the postsynaptic membranes. After annotations, the text files were imported into Matlab (MathWorks). The number and locations of vesicles, pits, endosomes were calculated using custom scripts (Watanabe et al., 2020). The example micrographs shown were adjusted in brightness and contrast to different degrees, rotated and cropped in Adobe Photoshop (v21.2.1) or Illustrator (v24.2.3). Raw images and additional examples are provided in Figshare.com/projects/dynamin_phase_separation. The macros and Matlab scripts are available at https://github.com/shigekiwatanabe/SynapsEM.

### Statistical analysis

Detailed statistical information was summarized in table S2. All EM data are pooled from multiple experiments after examined on a per-experiment basis (with all freezing on the same day); none of the pooled data show significant deviation from each replicate. Sample sizes were based on our prior flash-and-freeze experiments (∼2-3 independent cultures, over 200 images), not power analysis. An alpha was set at 0.05 for statistical hypothesis testing. Normality was determined by D’Agostino-Pearson omnibus test. Comparisons between two groups were performed using a two-tailed Welch two-sample t-test if parametric or Wilcoxon rank-sum test and Mann-Whiteney Test if nonparametric. For groups, full pairwise comparisons were performed using one-way analysis of variance (ANOVA) followed by Holm-Šídák’s multiple comparisons test if parametric or Kruskal-Wallis test followed by Dunn’s multiple comparisons test if nonparametric. All statistical analyses were performed and all graphs created in Graphpad Prism (v8).

All fluorescence microscopy data were first examined on a per-experiment basis. Sample sizes were ∼2-3 independent cultures, at least 50 synapses from 4 different neurons in each condition. An alpha was set at 0.05 for statistical hypothesis testing. The skewness was determined by Pearson’s skewness test in GraphPad Prism (v8). Since data were all nonparametric, Mann–Whitney test or Kruskal–Wallis test were used. p-values in multiple comparison were adjusted with Bonferroni correction. Confidence levels were shown in each graph. All statistical analyses were performed, and all graphs created in Graphpad Prism (v8).

### Theoretical calculations of dynamin nucleation and polymerization kinetics

We estimated the concentration of Dyn1xA in the cytosol based on fluorescence from endogenous proteins (Imoto et al., 2021) and also the data from a previous study (Wilhelm et al., 2014). The concentration is about 3.5 µM at a presynaptic terminal. The average volume of a synaptic terminal is 0.37 µm^3^ (Wilhelm et al., 2014). Based on the area of Dyn1xA puncta (0.1586 µm^2^), we calculated the volume of the condensate to be 0.05 µm^3^. From these values, the concentration of Dyn1xA in the puncta was estimated to be 11.6 µM, which is close the critical concentration for dynamin oligomerization (12.6 µM). At this concentration, the time that takes to recruit dynamin to the membrane becomes negligible since there are a sufficient number of molecules near the membrane to drive rapid polymerization. Based on the previous experiment (Roux et al., 2010), the polymerization speed is expected to be ∼ 361 nm/s with the concentration of 11.6 µM, assuming that dynamin polymerization if a first order reaction. Since dynamin pitch consists of 13 dimers of ∼ 10-14 nm (Chen et al., 2004; Cocucci et al., 2014), it will likely take 28 – 39 ms to form one dynamin pitch around the collar of endocytic pits when the liquid condensate is present.

When the concentration of dynamin is low, the critical number (*N*\*>1) of dynamin dimers must nucleate on the membrane before they can polymerize spontaneously. Hereby, the nucleation process is the rate-limiting step. According to the established theory (Lenz, 2009), the nucleation time can be determined analytically by relating the dynamin concentration (*C*), the critical number of dynamin dimer (*N*\*), and membrane mechanics. Consider the scenario with sufficient dynamin dimers in solution that are in chemical equilibrium with the membrane-bound dynamin molecules. According to this theory, the analytic formula of the nucleation time can be derived in the two extreme limits, and the real nucleation time will reside in-between. In the limit #1, where the membrane is soft and the dynamin polymer is stiff, the nucleation time, 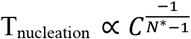. According to the above theory, in the limit #1, the nucleation time at 3.4 µM of dynamin will be ∼ 257 ms if the critical number of dynamin dimer, *N*\*, is 2 (*i*.*e*., dynamin tetrameter) (Ramachandran et al., 2007). In the limit #2, where the membrane is stiff and the dynamin polymer is soft, the nucleation time, T_nucleation_ is insensitive to the dynamin dimer concentration. The *in vitro* reconstitution experiments show that with 0.44 µM of dynamin in solution, the dynamin nucleation on a narrow membrane tubule (∼ 10-35 nm in radius) takes ∼ 2–5 seconds (Roux et al., 2010), which is the upper limit. Taken together, if the dynamin molecules do not form liquid droplets at a presynaptic terminal, the dynamin nucleation time alone is expected to range from ∼ 257 ms to several seconds. An additional ∼ 94–147 ms is necessary to polymerize dynamin around the collar once the nucleation initiates. Note that this estimated time is based on the optimal curvature-sensing condition that speeds up the membrane recruitment of dynamin. If the membrane curvature is less than optimal, then it will take even a longer time for dynamin to nucleate and polymerize on the membrane. Given that the ultrafast endocytosis completes within ∼ 50-100 ms (Watanabe et al., 2013a), the nucleation of freely-diffusing dynamin molecules may not be sufficiently fast to drive the ultrafast endocytosis.

## Source data availability

Source data are provided with the manuscript and through https://data.mendeley.com/datasets/bkn78y232y/draft?preview=1. All raw images are available upon request.

## Supplemental Information

