## Supplement figures and legends for "Dynamin forms liquid-like condensates at synapses to support ultrafast endocytosis"

Figure S1, Imoto, et al.

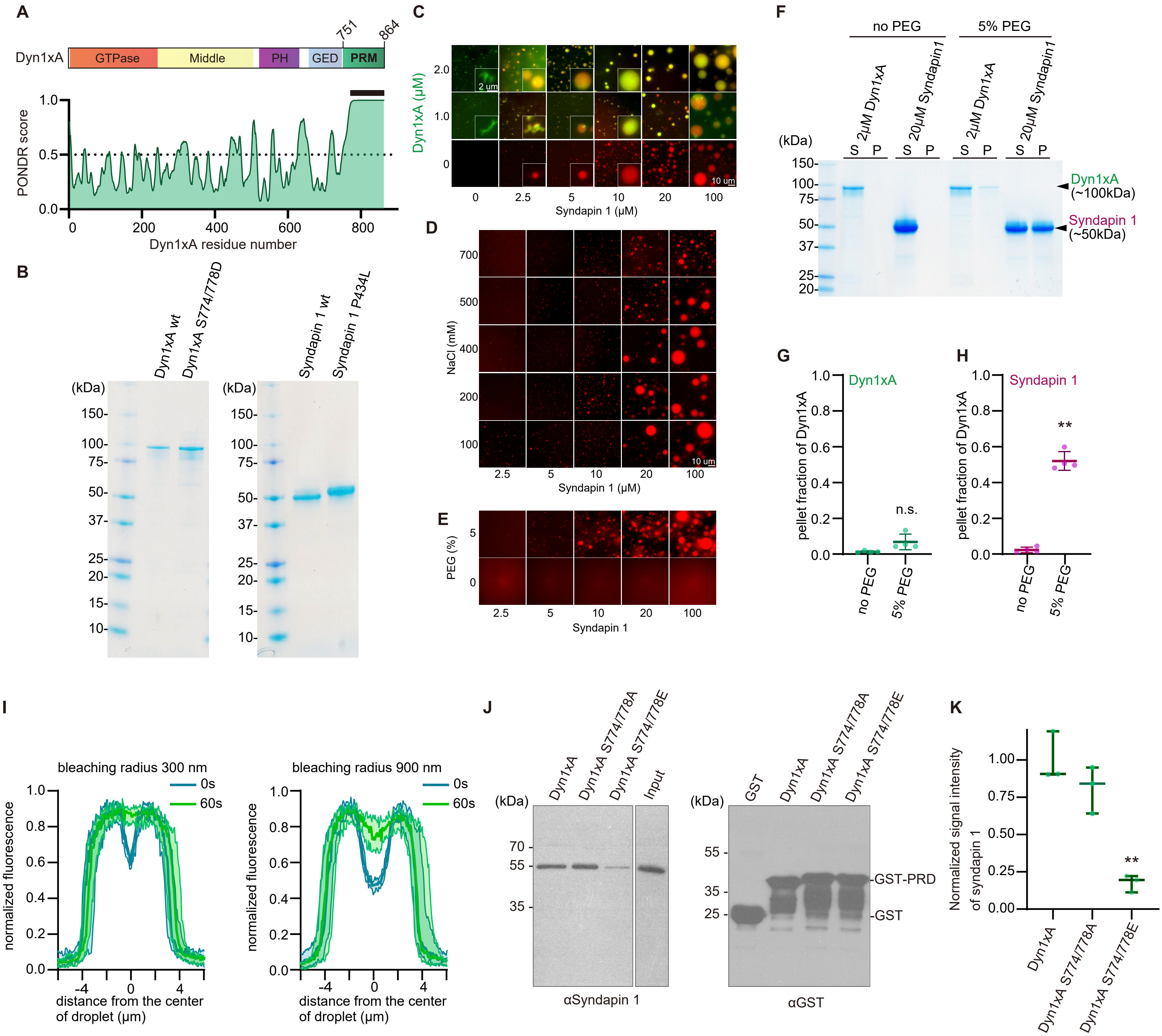

**Figure S1. Phase separation assay of Dyn1xA and Syndapin1 in *in vitro***

(A) Purities of recombinant proteins used in this study. Wild-type Dyn1xA (Dyn1xA wt), Dyn1xA S774/778D, wild-type Syndapin1 (Syndapin1 wt) and Syndapin1 P434L were purified from bacteria (see Material and Method) and assessed by SDS-PAGE and Gel code staining.

(B) Dyn1xA and Syndapin1 droplet formation at various protein concentrations. Enlarged images are shown in the insets.

(C) Syndapin1 droplets at various concentrations of protein and salt.

(D) Syndapin1 droplets with or without 5% PEG.

(E) Dyn1xA (2 μM) or Syndapin 1 (10 μM) were examined by sedimentation assay with or without 5 % PEG. The supernatant (S) and pellet (P) were collected by centrconfocalifugation at ×13,000 rpm for 10 min and then proteins were subjected to SDS−PAGE and Gel code staining.

(F, G) Pellet fraction of Dyn1xA (F) and Syndapin1 (G) calculated from protein bands in supernatants and pellet fractions in (E). *, $P < 0.05$. **, $P < 0.001$. Mean ± SEM are shown. n = 4 from two independent protein purification.

(H) Immunoblotting images showing a GST pull-down assay of the recombinant PRM of Dyn1xA, Dyn1xA S774/778A or Dyn1xA S774/778E and anti-Syndapin 1 antibody reactions. The assay was performed using the lysate of synaptosome fractions isolated from adult mouse whole brains. Right blot showing the blotting against anti-GST antibody used for normalization of signals in left blot.

(I) Normalized signal intensities of Syndapin 1 acquired from the immunoblotting using Dyn1xA PRM constructs shown in (H). Signal intensities were normalized to the amount of GST-PRMs The median and 95% confidence interval are shown.

N = 3 from different mouse brains in all cases. * $p < 0.05$. See Quantification and Statistical Analysis for the n values and detailed numbers.

Figure S2, Imoto, et al.

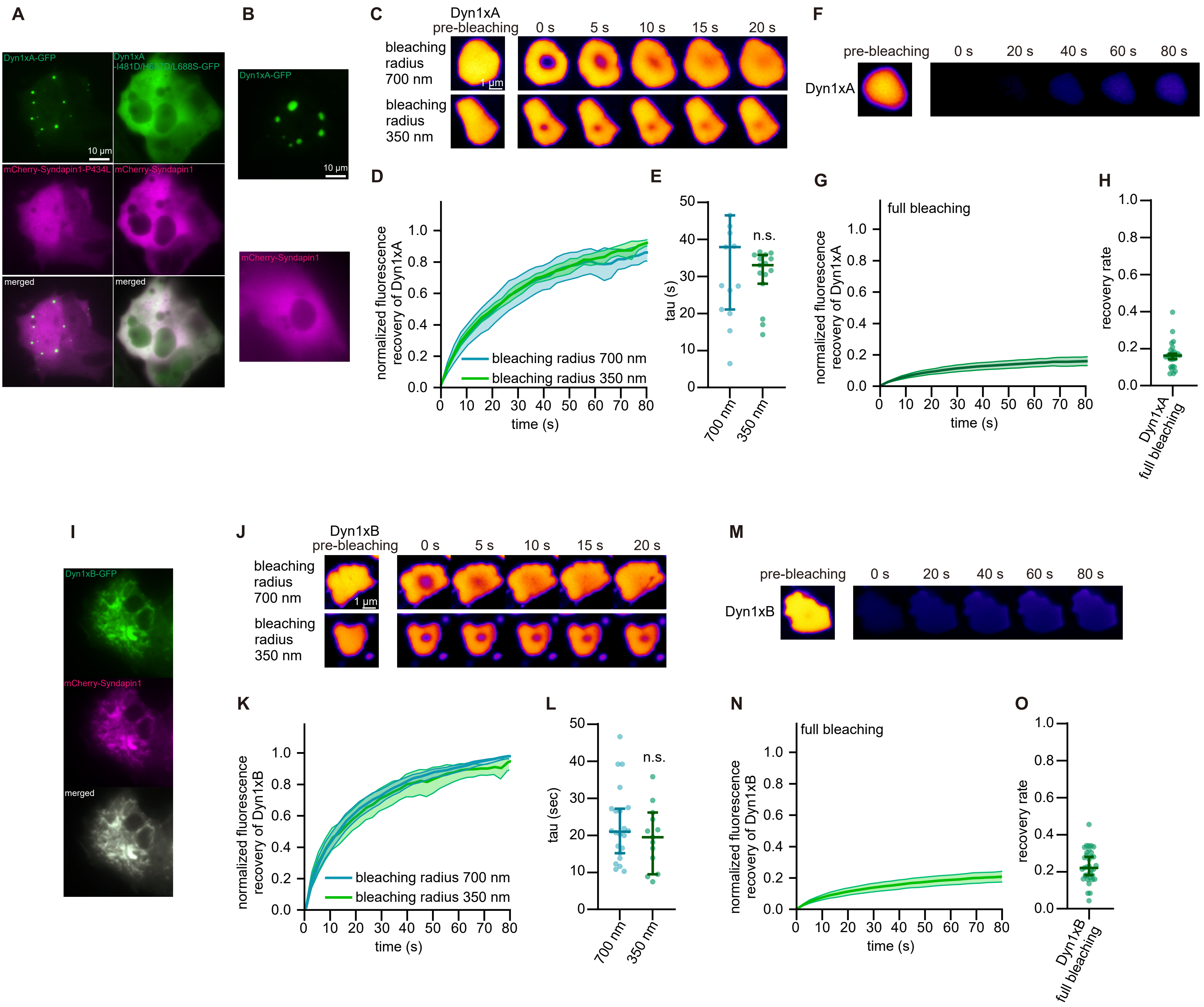

**Figure S2. Phase separation assay of Dyn1xA and Syndapin1 in COS-7 cells**

(A) Dyn1xA-GFP or Dyn1xA monomeric mutant (Dyn1xA-I481D/H637D/I688S)-GFP co-overexpressed in COS-7 cells with mCherry-Syndapin1 P434L or mCherry-Syndapin1 wild-type, respectively.

(B) Dyn1xA-GFP or mCherry-Syndapin1 expressed individually in COS-7 cells.

(C) Example time-lapse images of FRAP experiments inside of droplet-like Dyn1xA-GFP. Dyn1xA signals were photobleached at 480 nm using the ROI radius of 700 nm or 350 nm. Time indicates after the photobleaching.

(D) Normalized fluorescence recovery of Dyn1xA signals in different ROI sizes. Fluorescence signals were normalized between immediately (0 s) and 80 s after the photobleaching. Times indicate after the photobleaching. The median and 95% confidential interval are shown.

(E) The recovery time constant of Dyn1xA signals following the photobleaching using different ROI sizes in (D). The median and 95% confidence interval are shown. Each dot represents a droplet-like structure of Dyn1xA. $p < 0.0001$. Mann-Whitney test.

(F) Examples time-lapse images of FRAP experiments on the Dyn1xA-GFP droplet-like structure with the photobleaching laser covering the entire structure. Dyn1xA signals were photobleached at 480 nm.

(G) Normalized fluorescence recovery of Dyn1xA signals. Fluorescence signals were normalized at immediately (0 s) and 80 s after the photobleaching. Times indicate after the photobleaching. The median and 95% confidential interval are shown.

(H) The recovery rate of Dyn1xA signals at 80s after the photobleaching in (M). The median and 95% confidence interval are shown. Each dot represents a droplet-like structure of Dyn1xA-GFP.

(I) Dyn1xB-GFP and mCherry-Syndapin1 co-expressed in COS-7 cells.

(J) Examples time-lapse images of FRAP experiments on Dyn1xB-GFP and mCherry-Syndapin1 droplet-like structures. Dyn1xB signals were photobleached at 480 nm using the ROI radius of 700 nm or 350 nm. Time indicates after the photobleaching.

(K) Normalized fluorescence recovery of Dyn1xB signals inside the photobleached spots. Fluorescence signals were normalized between immediately (0 s) and 80 s after the photobleaching. Times indicate after the photobleaching. The median and 95% confidential interval are shown.

(L) The recovery time constant of Dyn1xB signals following the photobleaching using different ROI sizes. The median and 95% confidence interval are shown. Each dot represents a droplet-like structure of Dyn1xB-Syndapin1. n.s., no significance. Mann-Whitney test.

(M) Examples time-lapse images of FRAP experiments Dyn1xB-GFP and mCherry-Syndapin1 droplet-like structure of with the photobleaching laser covering the entire structure. Dyn1xB signals were photobleached at 480 nm.

(N) Normalized fluorescence recovery of Dyn1xB signals after the full bleaching. Fluorescence signals were normalized at just after (0 s) the photobleaching. Times indicate after the photobleaching. The median and 95% confidential interval are shown.

(O) The recovery rate of Dyn1xB signals at 80s after the photobleaching in (N). The median and 95% confidence interval are shown. Each dot represents a droplet-like structures of Dyn1xB-GFP and mCherry-Syndapin1.

n >15 droplets in more than 5 cells from 2 or 3 different cell preparations. See Quantification and Statistical Analysis for the n values and detailed numbers.

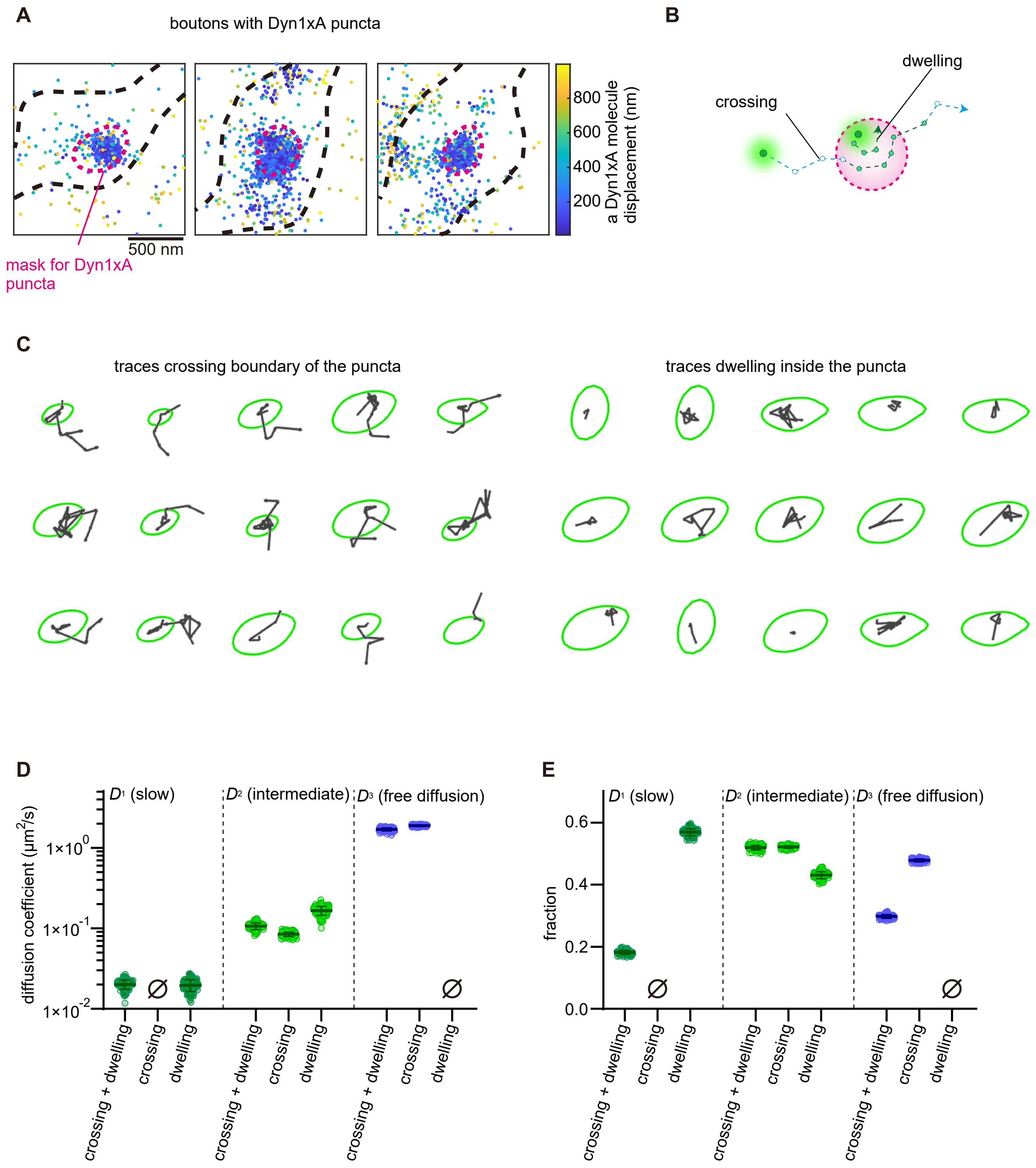
A
boutons with Dyn1xA puncta
500 nm
mask for Dyn1xA puncta
800
600
400
200
a Dyn1xA molecule displacement (nm)
B
crossing
dwelling
C
traces crossing boundary of the puncta
traces dwelling inside the puncta
D
diffusion coefficient (μm²/s)
1×10⁰
1×10⁻¹
1×10⁻²
D1 (slow)
D2 (intermediate)
D3 (free diffusion)
crossing + dwelling
crossing
dwelling
E
fraction
0.6
0.4
0.2
0.0
D1 (slow)
D2 (intermediate)
D3 (free diffusion)
crossing + dwelling
crossing
dwelling

**Figure S3. Phase separation assay of Dyn1xA puncta in neurons.**

(A) Example localization maps of photoactivated Dyn1xA molecules in bouton with the puncta. Color histogram indicates displacement of Dyn1xA molecues.

(B) Schematic showing traces of boundary crossing and staying inside (dwelling) which was used for calculation of diffusion coefficient.

(C) Example traces crossing boundary or dwelling inside of the puncta

(D) Diffusion coefficient of three components in Figure 3 L, M and N. n > 150 iterations.

(E) Fraction of diffusion coefficient of three components in Figure 3 L, M and N. n > 150 iterations.

Figure S4, Imoto, et al.

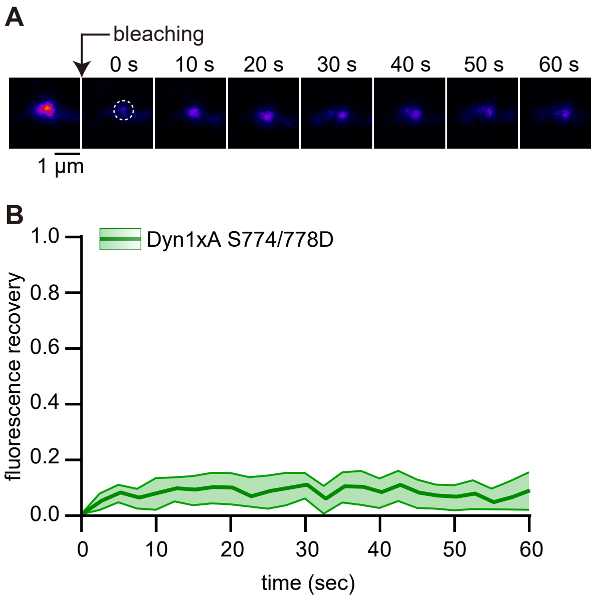

**Figure S4. Dephosphorylation of PRD regulates formation of Dyn1xA puncta**

(A) Examples live images of FRAP experiments of Dyn1xA-S774/778D-GFP puncta in presynapse. Dyn1xA-S774/778D-GFP signals were photobleached at 480 nm. Time indicates after the photobleaching.

(B) Normalized fluorescence recovery of Dyn1xA-S774/778D-GFP signals. Fluorescence signals were normalized to just after (0 s) photobleaching. Times indicate after the photobleaching. The median and 95% confidential interval are shown.
